# Unveiling Inter-Embryo Variability in Spindle Length over time: towards Quantitative Phenotype Analysis

**DOI:** 10.1101/2023.05.01.538870

**Authors:** Yann Le Cunff, Laurent Chesneau, Sylvain Pastezeur, Xavier Pinson, Nina Soler, Danielle Fairbrass, Benjamin Mercat, Ruddi Rodriguez Garcia, Zahraa Alayan, Ahmed Abdouni, Gary de Neidhardt, Valentin Costes, Mélodie Anjubault, Hélène Bouvrais, Christophe Héligon, Jacques Pécréaux

## Abstract

How to quantify inter-individual variability? When measuring many features per experiment/individual, this question becomes non-trivial. One challenge lies in choosing features to recapitulate high-dimensional data. This paper focuses on spindle elongation phenotypes to highlight how a data-driven approach can help tackle this challenge. We showed that only three typical elongation patterns describe spindle elongation in *C. elegans* one-cell embryo. We called them archetypes. These archetypes were automatically extracted from the experimental data using principal component analysis (PCA) rather than defined a priori. They accounted for more than 95% of inter-individual variability in a dataset of more than 1600 experiments across more than 100 different experimental conditions (RNAi, mutants, changes in temperature, etc.). The two first archetypes were consistent with standard measures in the field, namely the average spindle length and the spindle elongation rate in late metaphase and anaphase. However, our archetypes were not strictly corresponding to these classic, manually-set, features. The third archetype, accounting for 6% of the variability, was novel and corresponded to a transient spindle shortening in late metaphase. We revealed that it is part of spindle elongation dynamics in all conditions. It is reminiscent of the spindle elongation pattern observed upon kinetochore function defects. Interestingly, because these archetypes were all three present from metaphase on, it implied that spindle elongation around the anaphase onset is sufficient to predict its late anaphase length. We validated this idea using a machine-learning approach.

The inter-individual differences between embryos depleted from cell division-related proteins have the same underlying nature as inter-individual differences naturally arising between wild-type embryos. The same conclusion holds also when analysing embryos dividing at various temperatures. We thus propose that beyond the apparent complexity of the spindle and variability in the phenotypes of various gene depletions, only three independent mechanisms account for spindle elongation, weighted differently in the various conditions; meanwhile, no mechanism is specific to any condition. As such, given amounts of these three archetypes could represent a quantitative phenotype.

## 1 Introduction

While cell division is remarkably faithful, the mitotic spindle, key to ensuring a correct partitioning of the chromosomes, can take variable trajectories to achieve its task, varying its shape or organisation or its overall position in the cell [Fonseca et al., 2019, Farhadifar et al., 2016, Bouvrais et al., 2018]. Phenotype variation between genetically identical cells can arise from multiple causes, from random transcription rates to stochastic variations in internal chemical reactions [Snijder and Pelkmans, 2011, Niepel et al., 2013, Raj and van Oudenaarden, 2008]. This emergent variability can translate either into a gradation of phenotypes across cells, e.g. in expression levels of a fluorescent dye [Elowitz et al., 2002], or radically new cell behaviours, e.g. responsiveness to the induction of apoptosis [Spencer et al., 2009]. Threshold effects can induce the latter type of variability. Along that line, Raj et al. [Raj et al., 2010] investigated the development of the *C. elegans* intestine: high fluctuations in gene expression can lead to either viable or impaired intestinal development, even in a clonal population [Raj et al., 2010]. However, while this study exhibits the variability mechanism, it is generally challenging to identify whether a perturbation leads to quantitative or qualitative changes in phenotype. That is, whether one observes diverse grades of the same phenotypes or the appearance/disappearance of an entire phenotype, suggesting a switch. When it comes to the mitotic spindle, differences in protein quantities may cause variability, but it may also reveal a loosely constrained system [Doncic et al., 2006, Zhang et al., 2013, Barkai and Shilo, 2007, Montevil et al., 2016]. Thus variability in spindle trajectories was often viewed as noise, although it fosters the spindle’s ability to resist or adapt to internal defects like chromosome misattachment and external perturbations like changes in tissue environment [Shahrezaei and Swain, 2008, Knouse et al., 2018, Oegema et al., 2001, Itabashi et al., 2012, Bloomfield et al., 2020, Knouse et al., 2017, Heinrich et al., 2013]. Consistently, variability may increase cellular fitness in cancer [Sarkar et al., 2021]. Our paper proposes a methodology to characterise the nature of variability (qualitative versus quantitative) with a thorough quantitative analysis, using the well-studied and stereotypical cell division of *C. elegans* one-cell embryo as a representative example [Pintard and Bowerman, 2019].

To quantify cell-to-cell variability, one can choose a specific quantitative feature and display the variance over a given population, e.g. size of internal structures such as spindle or centrosomes [Farhadifar et al., 2015] or expression of a given gene of interest [Elowitz et al., 2002]. Studying variability in dynamical systems, like spindle kinematics during cell division, raises subsequent technical difficulties, namely measuring variability between trajectories. Disregarding the specific aspect investigated, mechanics or biochemistry, mathematical modelling can be instrumental [Tonn et al., 2019, Honegger and de Bivort, 2018, Acar et al., 2008]. The issue then shifts from comparing trajectories to comparing parameters from the fitting of the mathematical model to the experimental data, *id est* measuring the variance of each model parameter across a population of cells to capture the overall cell-cell variability. Although attractive, such an approach requires an established and integrative model, based on a priori assumptions, to recapitulate all the phenotypes observed experimentally.

Numerous models describe how wild-type cell divisions occur. However, because of the complexity resulting from both numbers and families of molecular actors involved, it is often required to focus on one part of cell division [Sönnichsen et al., 2005, Prosser and Pelletier, 2017, Guilloux and Gibeaux, 2020, Cai et al., 2018]. In particular, the correct partitioning of the chromosomes critically depends on the metaphasic spindle. Its functioning, especially the role of mitotic motors and microtubules, received much attention [Kapoor, 2017, Oriola et al., 2018, Elting et al., 2018, Oegema et al., 2001]. The spindle length is a classic and suitable entry point [Goshima and Scholey, 2010, Valfort et al., 2018]. It was already investigated through biological or modelling means [Needleman and Farhadifar, 2010, Wollman et al., 2008,Blackwell et al., 2017,Ward et al., 2014]. However, its mere observation left open many potential mechanisms: the complexity led to a broad range of non-consensual spindle elongation models, none recapitulating its dynamics fully. Moreover, tractable models usually rely on simplifying assumptions to capture the core principles of the biological system they represent. By its reductionist approach, modelling could overlook specific cell-cell differences that may account for unusual phenotypes.

In this paper, we present a complementary approach to model fitting. We classified spindle-elongation cell-cell variability using a data-centred approach, a common paradigm in biology [Greener et al., 2022]. From the spindle length over time measured during mitosis of the *C. elegans* one-cell embryo, we extracted Principal Component Analysis (PCA) projection as a blueprint of cell division. We investigated how each embryo may depart from this blueprint in genetically perturbed conditions to gain a phenotype taxonomy. We aimed to measure the variability observed across different embryos beyond the usual all-or-nothing classification and derived interpretable descriptors of such variability, aiming towards quantitative phenotypes. It is classically achieved by recapitulating phenotype through manually selected features. For instance, to study the genetic grounds of cell-division variability, Farhadifar and colleagues precomputed about 20 features, such as embryo size, division duration or centrosome oscillation duration and frequency, before applying a PCA [Farhadifar et al., 2015, Farhadifar et al., 2016]. In contrast, we propose a hypotheses-free approach, not using pre-computed features. We thus focused on data projection methods that reduce dimensionality, the most famous being PCA [Moon et al., 2019]. We expected the PCA to extract cell division’s key features and highlight biologically relevant mechanisms while discarding noise. Such an approach has been used to classify cell phenotypes according to gene expression or cell shapes [Park et al., 2014, Wang et al., 2020]. At the organism scale, it enabled describing nematode shapes and movements quantitatively across various strains and provided a list of descriptors to do so [Yemini et al., 2013]. We set to apply a similar approach to spindle dynamics during the division of the *C. elegans* one-cell embryo. Importantly, not enforcing a priori features allowed us to account for 95% of cell-to-cell heterogeneity of spindle elongation, with only three descriptors, compared to about 40% with about 20 pre-selected features [Farhadifar et al., 2016]. As we study trajectories over time, our automatically derived descriptors of spindle elongation can be seen as typical trajectories, called hereafter archetypes. Experimental elongations are then a linear combination of these archetypes added to the average elongation. Interestingly, our hypotheses-free descriptors partly meet the features largely adopted by the community, like the average spindle length.

## 2 Results

To understand the grounds of spindle behaviour variability, we derived descriptors of variability without a priori using PCA. Advantageously, it required no interpretative biophysical model. Key to such a data-centred approach was the choice of the dataset. We monitored the spindle elongation in the *C. elegans* one-cell embryo at typically 30 frames per second. We tracked the spindle poles with a 10 nm accuracy relying on our previously published automatic tracking methods [Pecreaux et al., 2006a, Pécréaux et al., 2016]. We analysed a set of published and unpublished experiments from the lab, corresponding to 1618 experiments covering 78 gene depletions or mutations (see table S1), and realised at temperatures ranging from 15°C to 25°C, referred to hereafter as the whole dataset. These experiments included non-treated conditions at various temperatures or labellings, and control conditions (L4440, see methods § 4.3). To further increase diversity, our dataset also comprised protein depletions through hypomorphic or penetrant RNAi, or mutants of genes involved in spindle positioning and mechanics. In particular, we tested all kinesins, that is, the 19 ones with interpro *Kinesin motor domain* homology [Blum et al., 2021, Siddiqui, 2002], microtubule-binding proteins (29 over the 64 with *microtubule-binding* GO term [Carbon et al., 2009, Srayko et al., 2005]. The experimental dataset also featured subunits *dli-1, dylt-1, dyci-1, dhc-1* from dynein complex and *dnc-1* from dynactin [Vaughan, 2012]. Overall, we targeted 52 genes with cell-division-variant phenotype or one of its descendants in the worm phenotype ontology [Sönnichsen et al., 2005, Kamath et al., 2003, Schindelman et al., 2011], out of 1191; we assumed that they were enough to challenge our approach as the dataset was diverse enough, combining various nontreated and genetic perturbations while requiring a reasonable dataset-acquisition effort to record embryo movies suitable for this study. Indeed, acquisitions had to be manually performed to reach the required accuracy.

### 2.1 Cell-cell variability in spindle elongation is recapitulated into three archetypes

#### Diversity of spindle elongation phenotypes

After filtering out the centrosome-tracking outliers (Methods 4.5), we computed the spindle length as the distance between the two centrosomes (Fig 1A-C). In the conditions tested in this article, we observed a variety of phenotypes departing from the average behaviour of the whole dataset, even restricting observations to embryos from the same strain and at an identical temperature (Fig 1D). Interestingly, a diversity of phenotypes was also visible among the non-treated condition alone, as exemplified by *γ*TUB::GFP strain (TH27) embryos in Fig 1E. The average spindle elongation for nontreated embryos (blue dashed curve) displayed twostep dynamics during metaphase and anaphase: A first mild increase in spindle length started about one hundred seconds before anaphase onset (referred hereafter as metaphase elongation), followed by a second quicker increase (referred hereafter as anaphase elongation), corresponding to the elongation in the literature. These two-step dynamics were observed in both the average elongation over the whole dataset and wild-type embryos. It is reminiscent of the “biphasic metaphase” observed by Goshima and Scholey [Goshima and Scholey, 2010]: in unperturbed condition, a quite constant spindle length in early metaphase followed by a rapid increase in late metaphase. When depleting proteins regulating microtubule stability and dynamics, trajectories could depart from the wild type, for instance targetting CLS-2 (Fig 1F). This CLASP protein stabilises microtubules, particularly at the kinetochore [Cheeseman et al., 2005, Srayko et al., 2005, Espiritu et al., 2012]. Upon partially depleting this protein through a *cls-2(RNAi)* treatment, we cancelled the metaphase elongation and rather yielded a clear spindle shortening before anaphase onset. It was consistent with the known defects in microtubule attachment at the kinetochore during metaphase [Lewellyn et al., 2010, Cheerambathur et al., 2017, Edwards et al., 2018]. The anaphase elongation also departed from the control to a variable extent: the three upper trajectories in Fig 1F (identified with an arrowhead) were typical of the so-called “spindle weakening” phenotype, resulting in a fast elongation of the spindle, much faster than the control one. From the mere pole-pole distance, it was impossible to state unambiguously whether it was happening by breaking the spindle because of the defective attachment of the microtubules at the kinetochore [Cheeseman et al., 2005] or during anaphase due to a weakened central spindle [Maton et al., 2015]. We arbitrarily assumed that fast elongation corresponds to anaphase onset in this condition. In a broader take, protein depletions caused a variety of phenotypes, raising two questions: (i) how to describe such a wide inter-individual variability beyond selecting a few features manually and taking the risk of missing key aspects of variability, and (ii) how to relate such a classification of variability to meaningful biological mechanisms.

**Figure 1:**
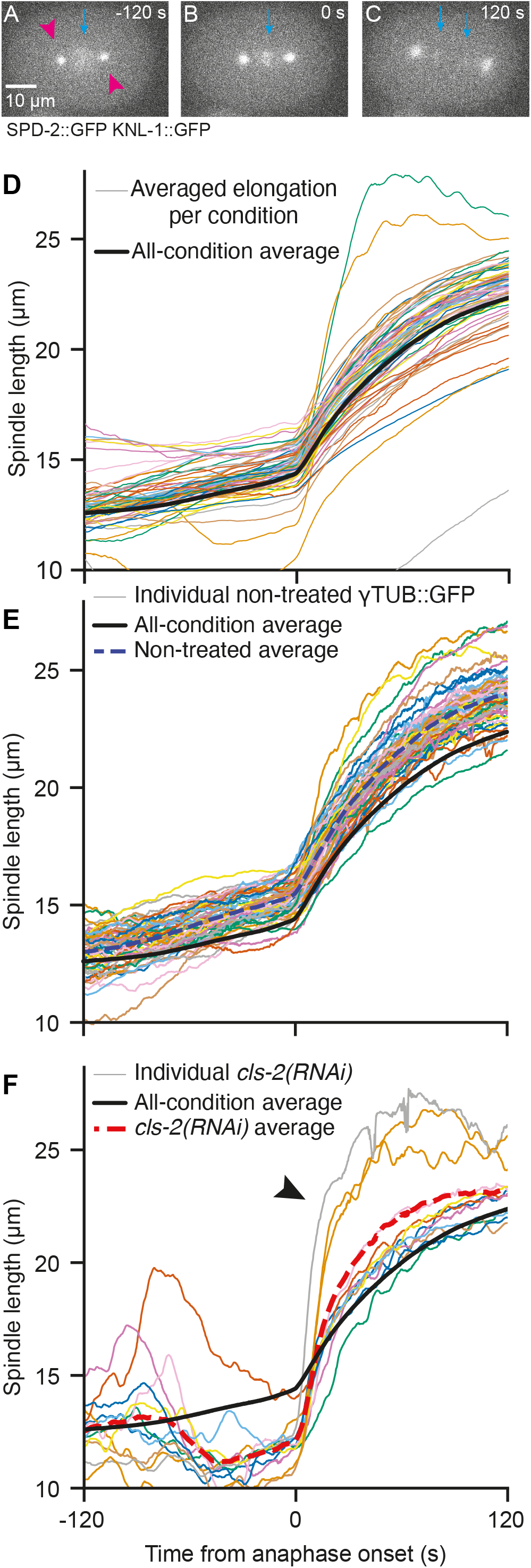
Diversity of the spindle length phenotypes at 18°C. (**A-C**) Exemplar stills of a typical embryo of strain JEP29, 120 s before, at anaphase onset and 120 s after. (magenta arrow-heads) Centrosomes were labelled through SPD-2::GFP and (cyan arrow) kinetochores through KNL-1::GFP. The scale bar represents 10 µm. (**D**) (thin coloured lines) Pole-pole distance (spindle length) averaged per condition and plotted during metaphase and anaphase for each of the 67 conditions, including 3 non-treated distinct strains, one control, and 63 gene depletions/mutations. All included embryos were imaged at 18°C (Table S1). Conditions with less than 6 embryos were not represented (see Suppl Table S1). Multiple conditions treating the same gene by RNAi or mutating it are merged. (**E**) The thin coloured lines depict the spindle lengths for individual non-treated TH27 embryos (*N* = 58), and the thick blue dashed line is the average of all embryos of this condition. (**F**) (thin lines) Spindle length for individual *cls-2(RNAi)* treated embryos (*N* = 12). The thick red dashed line corresponds to the average spindle length over these embryos. Arrowhead indicates the tracks showing fast spindle elongation revealing “spindle weakening”. In panels D-F, individual embryos and averaged tracks were smoothed using a 1.5 s-running-window median. The black thicker line corresponds to the average over the whole dataset, including all conditions.

#### Using projection methods to obtain a model-free description of variability

We set to use a linear and global method to ease the interpretability of the archetypes into biological features (see § 2.1 below) and accounted for the interplay between the various time stages of the division. We also aimed to perform an analysis without priors, so we focused on feature extraction and unsupervised algorithms [Jia et al., 2022]. We benchmarked several approaches and chose the one that maximised the within-condition to between-condition variability ratio (suppl. mat. § 3.1). The Principal Component Analysis (PCA) turned out to yield the second-best score (Table S4), while local linear embedding gave the best one. However, we used PCA as it was interpretable. Together with projecting trajectories, PCA produced eigenvectors, further termed archetypical elongation patterns, and allowed to compute their dosages in each embryo spindle-length curve, named below coefficients. Surprisingly, the first three descriptors of the PCA were necessary but also sufficient to provide most of the information needed to describe variability in our dataset, about 95% (Fig S1B). We further used three components, as it appeared to be the corner value of the L-curve, in an analogy to finding an optimum number of clusters in k-means algorithm or hierarchical clustering (Fig S1A) [Hastie et al., 2009]. For each elongation curve, the timepoint-wise difference from the whole-dataset average (Fig 1D, thick black curve) could be recapitulated into the weight given to each archetype (coefficient), i.e. the dosage of each of the three descriptors. Importantly, these descriptors (archetypes) were reminiscent of elongation patterns found in some perturbed conditions, either using RNAi or mutant (Fig 2A), suggesting they could have a biological interpretation. An exemplar reconstruction illustrates how three components were enough to account for a given single-cell spindle elongation (Fig S1C). Contrasting with previous studies based on pre-extracted features, we extracted the archetypes from the dataset rather than predefining them arbitrarily [Farhadifar et al., 2015, Farhadifar et al., 2016]. Our method thus provides hypothesis-free information regarding variability in spindle elongation.

**Figure 2:**
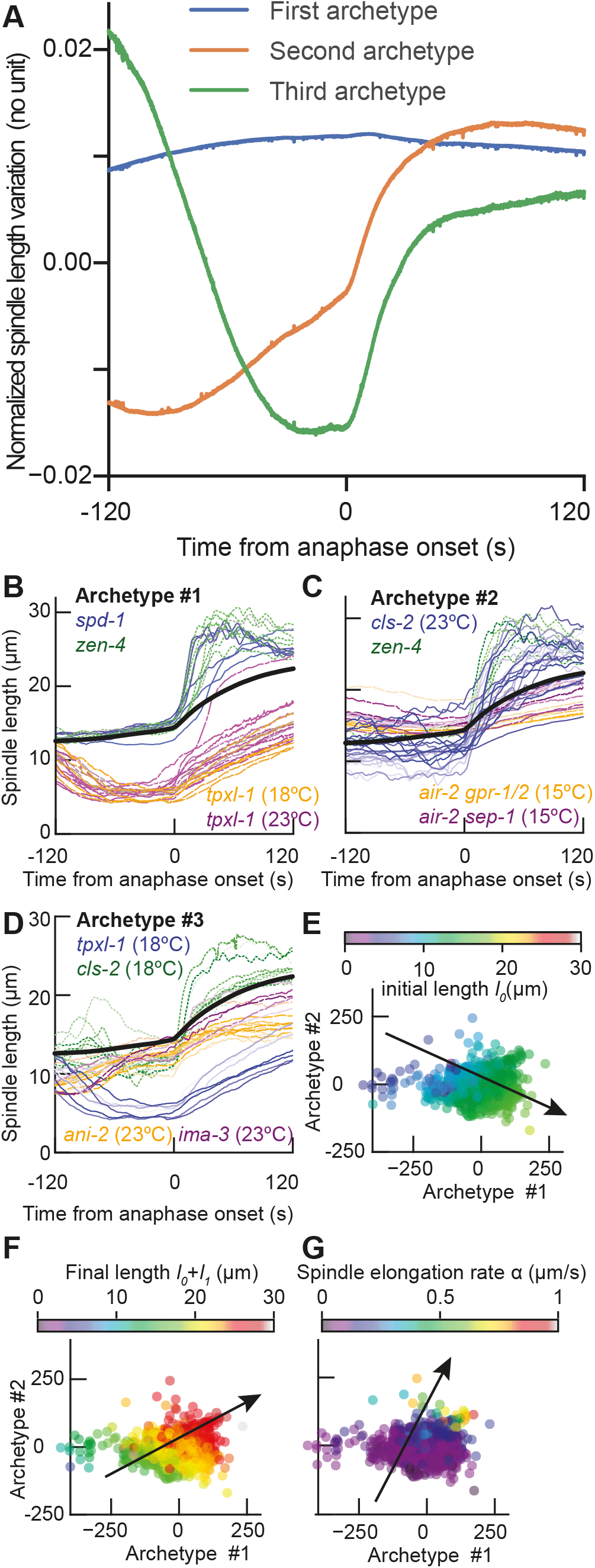
Principal component analysis of the spindle elongation and corresponding archetypes (eigenvectors) (**A**) The three main archetypes extracted by PCA over all conditions account for more than 95% of cell-cell variability in spindle elongation (see main text their plausible interpretation). (**B**) Spindle elongations from individual experiments for the two conditions resulting in the highest coefficients for archetype 1 averaged over the condition 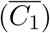, namely *γ*-TUB::GFP (TH27) embryos treated with *spd-1(RNAi)* 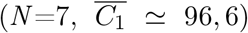 and *zen-4(RNAi)* 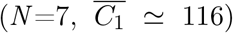. The lowest coefficients for archetype 1 were obtained by treating the same strain with *tpxl-1(RNAi)* and imaging at 23°C 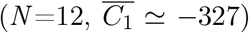 or 18°C 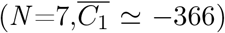 (**C**) Similar assay for archetype 2. Highest values were obtained with *γ*-TUB::GFP (TH27) embryos treated with *cls-2(RNAi)*, imaged at 23°C 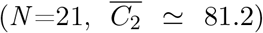 and *zen-4(RNAi)* imaged at 18°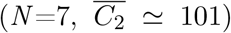. The lowest coefficients for archetype 2 were obtained treating the *air-2(or207)* mutant labelled with KNL-1::GFP and SPD-2::GFP (JEP31 strain) either with *sep-1(RNAi)* 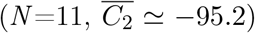 or *gpr-1/2(RNAi)* 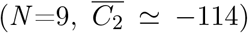, and imaging at 15°C. (**D**) Similar assay for archetype 3. Highest values were obtained with *γ*-TUB::GFP (TH27) embryos treated with *tpxl-1(RNAi)* 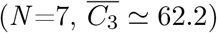 and *cls-2(RNAi)* 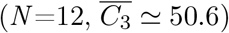 while lowest coefficients resulted from treating with *ima-3(RNAi)* 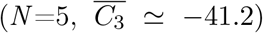 and *ani-2(RNAi)* 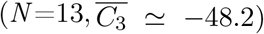. All considered embryos in panels (B-D) were imaged at 18°C except otherwise stated. (**E-G**) Fit of spindle elongation curves after [Farhadifar et al., 2015] and mapping the corresponding parameters on the PCA plane (see main text and Suppl Text § 1): (**E**) Average spindle length in early metaphase *l*_0_ and (**F**) in anaphase *l*_0_ + *l*_1_, and (**G**) elongation rate *α*. These three values are colour-coded, and an arrow depicts the axis along which these quantities vary (gradient) while we plotted the values of the two first coefficients of the PCA for embryos from the whole dataset except 16/1618 embryos whose cannot be fitted. 14 additional fits were excluded because of aberrant values.

#### Three main archetypes of spindle elongation

These main archetypes accurately describe the diversity in spindle elongation. The first archetype explained 70% of the overall variability in spindle elongation trajectories (blue curve, Fig 2A, S1B). Its roughly flat dynamics suggested that it accounted for the average spindle length. A higher corresponding coefficient reflects a shifted up spindle-length curve (Fig 2B). Interestingly, the average spindle-length was a feature used in many studies interested in the diversity of spindle elongation [Goshima and Scholey, 2010, Farhadifar et al., 2015, Dumont and Mitchison, 2009, Brown et al., 2007, Lacroix et al., 2018]. The second archetype explained 19% of the variability (red curve, Fig 2A, S1B) and comprised two elongation phases, during the metaphase (100 to 0 s from anaphase onset) and the anaphase (0 to 50 s). Therefore, the second archetype mainly captured the dynamics of spindle elongation: a high corresponding coefficient reflected a fast elongating spindle (Fig 2C). Such a feature was also previously identified as important to classify spindle elongation phenotypes, although only the anaphasic elongation was considered [Goshima and Scholey, 2010, Farhadifar et al., 2015, Scholey et al., 2016, Hara and Kimura, 2009]. The third archetype explained 6% of the variability (green curve, Fig 2A, S1B) and corresponded to a least an inflexion in the spindle elongation up to a plateauing that limited late-metaphase elongation and is observed in non-treated embryos, e.g. (Fig 2D, S2) [Pécréaux et al., 2016]. When more weighted, this archetype accounted for a spindle shortening, especially in *cls-2(RNAi)* (Fig 1F). Interestingly, targeting genes *ima-3* and *ani-2*, reported as reducing embryo length [Bouvrais et al., 2018], also produced extreme coefficients 3. We related that to improper metaphasic spindle length as embryo width was not proportionally reduced in these depletions, putatively misleading the spindle size regulation [Lacroix et al., 2018]. In a broader take, the phenotype of spindle-limiting-elongation up to shortening was previously reported upon misattachment of kinetochores by the microtubules [Lewellyn et al., 2010, Cheerambathur et al., 2017, Edwards et al., 2018] but not reported as a classifier of phenotypes in previous studies. Beyond proposing late metaphase shortening as a novel feature, we found objectively two archetypes reminiscent of known ones, namely the average spindle length in metaphase or at anaphase onset, and the elongation rate during early anaphase.

In a reversed perspective, we compared the three archetypes to arbitrarily chosen features used in previous studies to better understand the corresponding biological mechanisms. We fitted individual embryo elongation curves using non-linear least-squares to extract three features: the average spindle length at early metaphase, the final spindle length in anaphase, and the elongation rate (Supplemental text § 1). We then mapped the above manual features on the two first PCA dimensions (Fig 2E-G) (Supplemental method § 3.2). We found that the initial length is mostly related to our first coefficient, as depicted by the black arrow (Fig 2E). Meanwhile, the spindle’s final length combined coefficients 1 and 2. Finally, we found a good alignment of the elongation rate with our second coefficient of the PCA, as expected. In a broader take and beyond proposing archetypes, we found that only three components (archetypes) are needed to recapitulate most of the spindle elongation kinematics, suggesting that few mechanisms underlie the complexity of spindle elongation phenotypes.

### 2.2 Anaphase onset as a turning point

Because the two first archetypes, which spanned over metaphase and anaphase, accounted for 89% of the variability, we asked whether the spindle length in metaphase could correlate with the anaphasic one. We computed the spindle length in late anaphase (*l*_*LA*_) as the average of the 300 last data points spanning between 111.7 s and 120 s after anaphase onset. Notably, beyond being a marker of correct spindle functioning, this feature correlates with spindle final position and, therefore, cytokinetic furrow positioning [White and Glotzer, 2012, Rappaport, 1971, Knoblich, 2010, Bringmann and Hyman, 2005, Pintard and Bowerman, 2019]. We used a machine learning approach to investigate whether late anaphase spindle length was set early during division (Suppl Text §3.3). Spindle length during a 25 s interval starting 5 s before anaphase onset provided an accurate prediction of length minutes later (purple and green shadings, respectively, Fig 3A). Comparing predicted spindle late-anaphase lengths and experimental ones, we obtained a Pearson coefficient *R* = 0.82 with *p <* 10^*−*15^ (Fig 3B). We next wondered how this correlation depended on the starting time of the 25 s interval. Considering Pearson R as the *predictive power*, we computed it when sliding the interval (Fig 3C). We observed a steep increase in the predictive power around anaphase onset. In other words, most late-anaphase spindle length variability depends on the mechanisms already present at anaphase onset.

**Figure 3:**
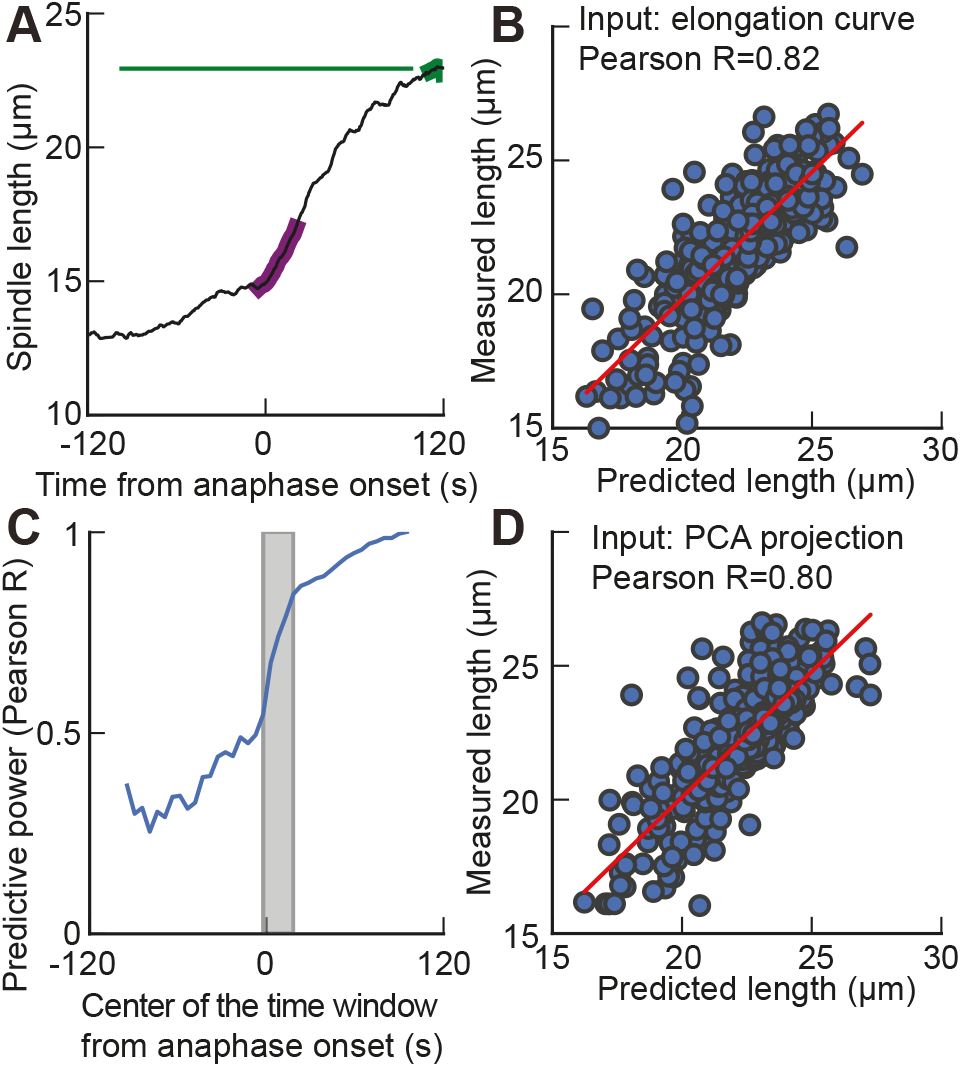
Machine learning (ML) predicts the spindle late-anaphase length *l*_*LA*_. (**A**) (black line) elongation of an exemplar non-treated *γ*-TUB::GFP embryo, imaged at 18°C, highlighting (purple) the (*−*5, 20) s interval used as input of algorithm and (green) the (111.7, 120) s interval from anaphase-onset to compute the average spindle length at late-anaphase. (**B**) Using embryos in the testing set (32%, i.e. 576), we plotted the measured spindle late-anaphase length versus the predicted final length and obtained a high correlation (R=0.82, *p <* 10^*−*15^) when inputting an elongation curve over the (−5, 20) s interval as input to the ML network. (**C**) We slid this 25 s interval by 5 s starting at *−*120 s and ending at 120 s and computed the Pearson coefficient, as above, denoted predictive power. The grey shading region corresponds to the interval used in other panels. (**D**) Using embryos in the testing set, we plotted the measured spindle late-anaphase length versus the predicted final length and obtained a high correlation (R=0.80, *p <* 10^*−*15^) when inputting the PCA projection (coefficient) computed on the elongation curve over the (−5, 20) s interval as input to the ML network. Details about the algorithmic approach is provided in Suppl Text §3.3.

We next asked whether this information announcing the late-anaphase spindle length was captured by PCA projection. We thus trimmed the elongation curves to the (−5, 20) s interval, did the PCA projection as described above and kept only three components (archetypes). We then performed a similar machine learning assay (Suppl Text §3.3). We obtained a Pearson coefficient *R* = 0.80 with *p <* 10^*−*15^ (Fig 3D) when comparing predicted and measured late-anaphase lengths. We concluded that the spindle characteristics supporting the prediction of the late-anaphase length were well recapitulated by the coefficients of the three archetypes, at least in late metaphase and early anaphase.

### 2.3 Robustness of the main archetypes

#### Robustness to dataset composition

The main archetypes are automatically extracted from the dataset to describe its heterogeneity best. Therefore, one can wonder whether the rather extreme phenotypes corresponding to high coefficients for archetypes are defining the archetypes alone (Fig 2B-D). In other words, firstly, are milder phenotypes well represented by the proposed three archetypes? and secondly, how robust are these archetypes with respect to the composition of the dataset? To investigate this issue, we first performed a bootstrap sampling of the experimental dataset. We randomly selected 500 experiments among the 1618 ones of the dataset and repeatedly computed the archetypes (eigenvectors of the PCA) (methods § 3.4). We found mild changes in archetypes despite considering only 31% of the dataset (Fig 4). It suggests that our archetypes are present in most of our embryos rather than set by a few extreme phenotypes.

**Figure 4:**
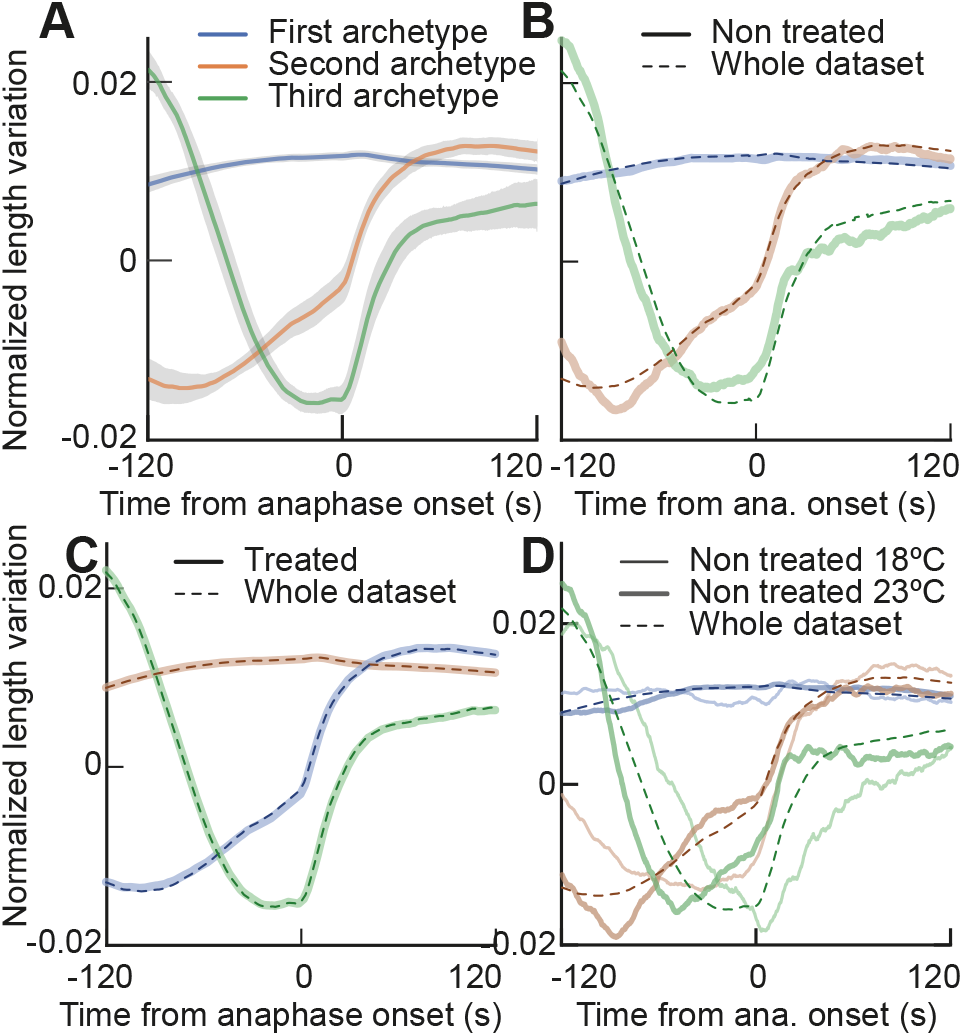
Variations of the archetypes upon dataset changes. (**A**) (lines) Average of the three first PCA archetypes (eigenvectors) *±* (shading) two times their standard deviations, computed over a 500 embryos subset of the data (disregarding conditions). We repeated this bootstrapping 500 times to obtain standard deviations. (**B**) (thick lines) Three first PCA archetypes were computed considering only the non-treated conditions (*N* =129) and compared to (dashed lines) archetypes extracted from the whole set of conditions (*N* =1618). (**C**) (thick lines) Three first PCA archetypes computed considering only the treated conditions (*N* =1308), meaning RNAi or mutant without L4440 controls, and compared to (dashed lines) archetypes extracted from the whole set of conditions. **D**) Three first PCA archetypes computed considering only the non-treated conditions at (thick lines) 23°C (*N* =71) and (thin lines) 18°C (*N* =58), compared to (dashed lines) archetypes extracted from the whole set of conditions. In all panels, the elongation curves were smoothed with a 1.5 s running-median filtering before computing PCA. Explained variances are reported in Table S5

Going further, we focused on the sole non-treated embryos. These embryos demonstrate a spindle elongation close to the average behaviour, that is, the (0,0) position in the PCA plan (as shown in figure S3). Considering only these embryos can be seen as a more extreme version of bootstrap, in which most extreme phenotypes are indeed removed. Strikingly, we found similar archetypes (Fig 4B). We also tested the use of only the treated embryos (excluding the control experiments using L4440 RNAi) and again found similar archetypes (Fig 4C).

Therefore, disregarding the subset of the dataset, the same overall interpretations remain: the first archetype accounts for average spindle length, the second for spindle elongation and the third for a spindle temporary limited-elongation / shortening. While a shortening was not visible at first sight in most non-treated embryos during late metaphase (Fig 1E), the PCA revealed that this phenomenon is still critical to explain cell-to-cell heterogeneity in these embryos (Fig S1BCE, Suppl Table S5). More generally, this suggests that the genetic perturbations explored in this study, while covering a broad range of documented molecular actors involved in spindle elongation, can be accounted for by changing the weights of the archetypes but do not require archetype changes. That is no qualitative change in the spindle elongation phenotype is observed upon perturbations. Moreover, the contributions of each archetype to describe the overall variability of nontreated embryos are comparable to those obtained with the whole dataset (Suppl Table S5).

#### Robustness to experimental conditions

Variability can also arise from changes in the environment to which the organisms might respond. Worms in their natural environment experience various kinds of stress, among which changes in the temperature have received attention [Pecreaux et al., 2006a, Begasse et al., 2015]. We wondered whether variability in spindle elongation would have the same underlying archetypes at two different temperatures since they correspond to two different spindle elongation phenotypes (Fig S4). We repeated the PCA projection considering two sets of non-treated embryos at 18°C and 23°C, and observed that the general shape of the archetypes are similar across temperatures and compared to the ones extracted from the whole dataset (Fig 4D). Importantly, as expected, the archetypes appeared horizontally squeezed at the higher temperature because the division pace is increased at higher temperature [Begasse et al., 2015]. It likely reflects a higher dynamics of molecular components, like the microtubules and molecular motors, although all are not identically sensitive to temperature changes [Chaaban et al., 2018, Kushwaha and Peterman, 2020, Yadav and Kunwar, 2021]; for instance, it leads to faster anaphase spindle rocking at higher temperature [Pecreaux et al., 2006a]. It was also noteworthy that the second archetype did display a first elongation in late metaphase, not visible in the 18°C counterpart. We also investigated the contri-bution of each archetype to explain variance and observed variability (Suppl Table S5). Genetically perturbed embryos, with some extreme phenotypes, relied a bit more on archetypes 2, mildly increasing their corresponding explained variance. Finally, and as a control, we selected the embryos from the functional group *kt* as they display a very peculiar elongation pattern (Table S1, Fig S6B). Not surprisingly, PCA applied to these embryos resulted in different components (Fig S6). We concluded that provided that a variety of elongation patterns is fed into the PCA through either a large enough set of non-treated embryos or a variegated set of protein depletions, the archetypes’ general shapes were conserved across temperatures, a panel of genetically perturbed embryos versus non-treated or control ones, which suggested that while dynamics of elongation are influenced by these changes, the nature of variability did not change.

### 2.4 From archetypes to phenotypes

#### Interplay between the spindle elongation phenotype and the projection

We used partial RNAi-mediated protein depletion in many cases (Suppl Table S1). We reckoned that the shift of PCA coefficients from control may depend on the penetrance of the RNAi. Providing a general demonstration of such a link would be out of the scope of this paper. We instead offered an example: we varied the amount of a known microtubule-associated protein, the depolymerising kinesin KLP-7. Its depletion caused decreased growth and shrinkage rates, and reduced rescue and catastrophe rates [Srayko et al., 2005, Lacroix et al., 2014], causing microtubule-chromosome attachment defects [Wordeman et al., 2007, Ems-McClung and Walczak, 2010]. It is reported to cause only mild spindle length defect upon hypomorphic treatment (Fig 5A), while spindle breakage happens only upon penetrant one [Greenan et al., 2010,Grill et al., 2001]. It made KLP-7 an excellent candidate to test the link between penetrance and PCA coefficients. Thus, we investigated embryos where this protein was fused with mNeonGreen at the locus and depleted by RNAi. We observed a larger first coefficient in correlation with a higher amount of fluorescence, i.e. depletion causes a lower “dose” of archetype 1 (Fig 5B). It corresponded to a Spearman correlation coefficient between the first coefficient and the fluorescence of *ρ* = 0.35 (*p* = 0.016, *N* = 36). Because of the low number of samples, we used a permutation test to show a mild significance of this correlation. The second and third coefficients did not appear significantly correlated with the level of fluorescence. In the case of KLP-7, the variability of the phenotype, quantified by our PCA coefficients, reflects the grading of depleting by RNAi. Conversely, it suggests that variability of penetrance likely results in spreading the coefficient values within a given condition cloud. Other causes of spreading are likely present. Indeed, the non-treated condition also inherently displays variability (Suppl Fig S3, Suppl File 2) corresponding to the distribution of elongation curves (Fig 1E).

**Figure 5:**
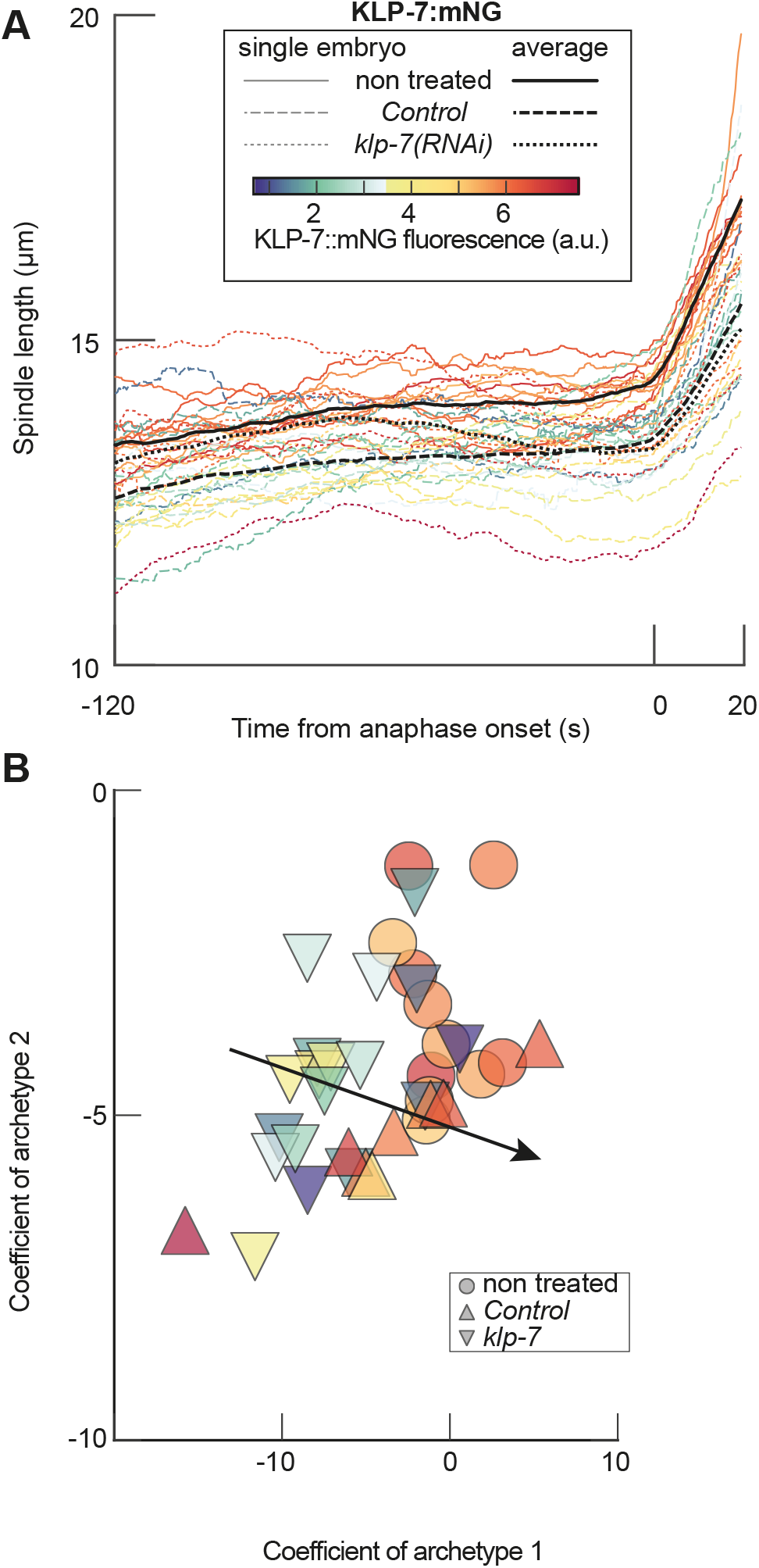
PCA coefficients depend on the penetrance of RNAi. (**A**) Spindle elongation trajectories for the depletion of KLP-7^MCAK^in KLP-7::mNG background (strain LP447) in three conditions: (dotted lines) *N* =18 *klp-7(RNAi))* treated embryos; (dashed lines) *N* =8 control embryos (L4440 treated); and (plain lines) *N* =11 non-treated embryos. The thick lines report the averages of each condition. (**B**) Projection of these conditions on the PCA established on the whole dataset. We plotted the coefficients corresponding to the two first archetypes. The black arrow depicts the axis along which this fluorescence varies (gradient), computed similarly to Fig 2EFG. The three conditions reported here were not included in the initial dataset used to generate PCA archetypes. Acquisitions were performed at 18°C. The line or marker colour encodes the fluorescence level of KLP-7::mNG (Methods §4.6).

#### From spindle elongation pattern to gene function

Having established that the position of an experiment in the space of PCA coefficients is a robust and *bona fide* representation of its phenotype of elongation, we reckoned that the median position over all replicas of the same condition may map the phenotype in the PCA plane by linking gene functions and the weights of the archetypes. We computed the median coefficient for each condition and mapped them in the PCA plane. Since gene ontology terms or worm phenotype ontology turned out to be too generic, we tagged each condition manually with the main function of its corresponding protein (group column in Suppl Table S1) during embryonic division. Plotting the corresponding groups in the PCA coefficient space indicated partially overlapping clusters (Fig 6, Suppl File 1). To test to which extent each coefficient supported this grouping, we used the Kruskal-Wallis test for each coefficient. While coefficient 1 did not really support clustering (*p* = 0.12), coefficients 2 and 3 are discriminative (*p* = 0.0044 and *p* = 7.34 *×* 10^*−*5^, respectively). We shuffled the group labels and computed the Kruskal-Wallis H values; we repeated this 10000 times, and the corresponding distribution supported our conclusions (Suppl Fig S5). We concluded that our PCA analysis could be instrumental in suggesting gene function by looking at their neighbours in the PCA space.

**Figure 6:**
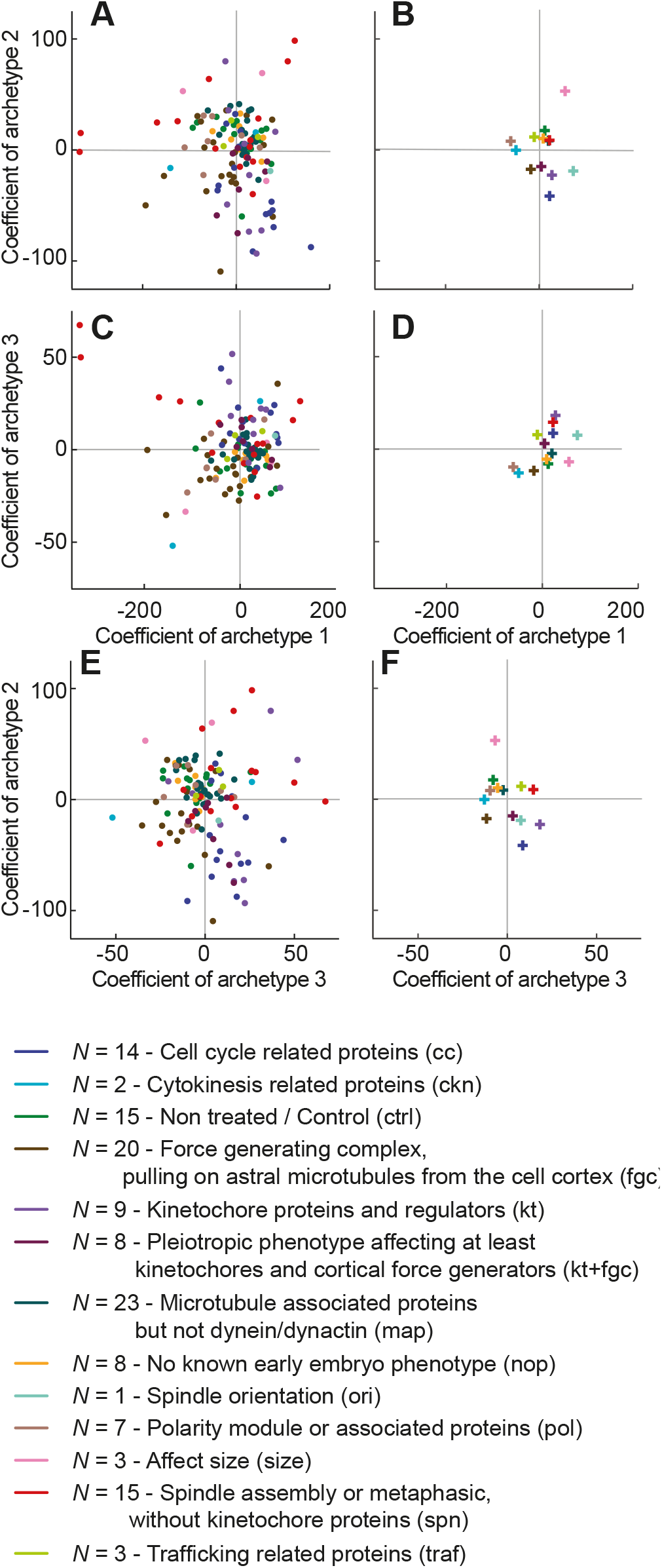
PCA coefficients as a quantitative phenotype. Each experiment is projected by PCA, and then a median is computed per condition. The resulting scatter plot is attached as an interactive plot (Suppl File 1). (**A, C, E**) report on the coefficients 1 and 2, 2 and 3, 1 and 3 respectively. We colour-coded the conditions depending on their functional group as reported in (Suppl Table S1). (**B, D, F**) depict the corresponding coefficients for each group computed as the median of the values per conditions plotted on the left-hand side panels. The corresponding spindle elongations are reported at Suppl Fig S7. The colour code for the group is shown in the bottom part, together with the abbreviated group within parentheses.

## 3 Discussion

The specific question of extracting quantitative descriptors (in this study, the archetypes) of inter-individual variability has received interest from the cellular level [Farhadifar et al., 2015, Valfort et al., 2018] to the individual level [Cronin et al., 2005, Gyenes and Brown, 2016, Yemini et al., 2013] in the C.*elegans* research community. Our work complements and departs from these studies in several ways. In their study, Farhadifar et al. aimed at linking phenotypic diversity in spindle elongation with mutation accumulation over several generations or with evolutionary divergences between various strains [Farhadifar et al., 2015]. In contrast, rather than tackling evolutionary genetics, we focus here on linking spindle elongation phenotypes with mild perturbations of key molecular players in the most standard C.*elegans* lab strains to shed light on the molecular mechanisms underlying variability in spindle elongation. Moreover, we did not use preselected features to describe spindle elongation but instead, let the projection method extract the most suited descriptors of variability for our dataset. It enabled us to uncover the third archetype, displaying a limited elongation up to shortening during late metaphase, present in all conditions, including the non-genetically perturbed ones.

Interestingly using PCA directly on time-series provides interpretable descriptors of variability (archetypes). Our first two archetypes, reminiscent of the spindle-length average and elongation rate, respectively, correlate with intuitively-set features (Supplemental text § 1, Fig 2E-G) [Farhadifar et al., 2015]. However, extracting the archetypes without a priori and making them independent allowed us to focus on more subtle aspects. For instance, the second axis of the PCA, that is, the “dose” of the second archetype, did not perfectly align with the spindle elongation rate measured during early anaphase, albeit being correlated with it (Fig 2F). It suggests that a slightly different descriptor is more accurate in describing the dynamics of spindle elongation. The second archetype displays two different slopes (Fig 2A, orange curve), one for the late-metaphase elongation – roughly starting 100 seconds before the metaphase-anaphase transition – and one for the anaphase elongation – starting at metaphase-anaphase transition. Our second archetype relies on both these slopes and not only the second one, starting at anaphase onset. This hints at mechanisms in spindle dynamics which would be conserved before and after anaphase onset. Consistently, we found that spindle length around anaphase onset is a predictor of spindle length in late anaphase meanwhile considering only late metaphase offers already predictions that reasonably correlate with actual final lengths (Fig 3).

Even more surprising, the third descriptor identified by this study highlights a transient spindle limited-elongation phenotype during late metaphase. A plain shortening has been described in particular conditions with defective kinetochore-microtubule attachment dynamics [Ozlu et al., 2005, Lewellyn et al., 2010, Cheerambathur et al., 2013, Edwards et al., 2018, Cheeseman et al., 2005]. The corresponding conditions present in this study displayed a significantly increased third coefficient (Suppl Text § 2, Fig S3, Table S6). Yet, it has never been considered as a widespread phenomenon, which is instrumental in describing variability in spindle elongation in all conditions. At first glance, it might indeed seem paradoxical to include such a spindle-shortening phenotype observed in only some specific experimental conditions. Indeed, this coefficient contributes to accounting for variance even for non-treated embryos and many conditions, even not in the kinetochore-related groups have a third coefficient departing from zero (Suppl Table S5, Fig S3, Fig 6). Yet, this phenomenon *naturally* occurs in wild-type embryos, but on a smaller magnitude than in genetically perturbed ones.

This statement raises the fundamental question of defining what is a phenotype in our biological system. Indeed, our archetypes can be described as phenotypes for extreme embryos (Fig 2B-D), complying with the usual definition of the term: observable characteristics. Previous papers have indeed reported shorter spindles and spindle weakening leading to fast elongation or spindle shortening [Goshima and Scholey, 2010, Wuhr et al., 2009, Barisic et al., 2021, Edelmaier et al., 2020, Scholey et al., 2016,Kapoor, 2017]. Yet, we show that the non-treated embryo phenotype is a mixture of all three archetypes. It suggests that rather than describing an experiment with discrete descriptors, that is, whether a phenotype is absent or present, it would be more accurate to adopt a continuous description, namely the relative weights of archetypes. In a broader take, such a continuous description of experiments could lead to more straightforward interpretations. Indeed, a phenotype would most likely not disappear completely when mildly changing experimental conditions but would diminish in intensity. Along that line, we mapped our different experiments using the proposed PCA to determine closely related genes through their quantitative phenotypes (Fig 6).

Using a dimensionality reduction approach without a priori also enabled us to investigate the complex spindle and propose only three descriptors [Kapoor, 2017, Oriola et al., 2018, Elting et al., 2018]. Thus, the variability can be described by a small number of descriptors. It is likely that, in many cases, the use of data-driven descriptors could reduce the number of descriptors from hundreds or thousands to a handful. For instance, quantifying C.*elegans* shape and movements is difficult, given the large apparent diversity in a given population. While previous studies have undergone a meticulous study of variability between individual shapes and behaviours by selecting up to several hundreds of features, unsupervised projection methods have identified four archetypes which recapitulate 97% of the observed variability in the dataset [Gyenes and Brown, 2016, Stephens et al., 2008]. Because variability relies on a small number of descriptors in these both very different cases, it suggests that beyond the apparent complexity, only a few independent mechanisms may be at work.

A small number of variability descriptors, spanning across metaphase and anaphase, can also be understood as strong constraints for the biological system studied. As experimental data can be rebuilt only using these few descriptors, it follows that the range of possible experimental observations is bounded. For instance, the second archetype shows a first pre-anaphase elongation as well as anaphase elongation at anaphase onset. It means that a given experiment with a high coefficient for the second archetype would then have both a first and an anaphase elongation faster than average. This link between spindle elongation dynamics occurring before and after sister chromatids separation is in itself interesting to highlight. Indeed, the establishment of the central spindle, although not fully understood, requires a re-assembly from scratch of the microtubules [Nahaboo et al., 2015, Maton et al., 2015, Laband et al., 2017, Yu et al., 2019,Khmelinskii and Schiebel, 2008,Scholey et al., 2016]. While the central spindle only appears after anaphase onset, we found a clear correlation between spindle-elongation dynamics before and after chromatid separation (Fig 3). This highlights that our approach not only quantitatively detects variability but also reveals robust aspects of spindle elongation.

Finally, mapping conditions in the PCA plane (Fig 6) provides indications of similarity between conditions. Such similar displacements along one of the axes in the PCA map often indicate some commonality in the underlying molecular mechanisms. As such, when investigating a spindle-related gene with little to no documented function, one could partially impair its expression, monitor spindle elongation and project the result in the PCA map. Therefore, our PCA analysis could turn into a prospective tool for finding gene candidates for a mechanism by similarity to known players. This approach could also be instrumental in comparing various strains of C.*elegans* or even various nematode species. Indeed, distance in the PCA plane can be seen as a phenotypic distance between conditions [Xu et al., 2013, Sheehan et al., 2008,Mathur and Dinakarpandian, 2012,Kulmanov et al., 2021, Gan et al., 2013], more quantitative than the ones using phenotype ontology [Schindelman et al., 2011].

## 4 Material and Methods

### 4.1 Culturing *C. elegans*

This study partly re-used experiments, which we have previously published, as referenced in Table S1. In all cases, *C. elegans* nematodes were cultured as described in [Brenner, 1974] and dissected to obtain embryos. The strains were maintained and imaged at temperatures between 16°C and 25°C (Table S1). The strains were handled on nematode growth medium (NGM) plates and fed with OP50 *E. coli* bacteria.

### 4.2 Strains used in this study

Strains carrying mutations or fluorescent labels in use in this study are detailed in Table S2. Some strains were obtained by crossing, as detailed in this table.

### 4.3 Gene inactivation through protein depletion by RNAi feeding

RNA interference (RNAi) experiments were performed by feeding using either Ahringer-Source-BioScience library [Kamath et al., 2003] with the bacterial strains detailed in Suppl Table S1, either bacteria transformed to express dsRNA targeting the desired gene. The bacterial clone (bact-16) targeting *par-4* is a kind gift from Anne Pacquelet and was, in turn, obtained from [Lee et al., 2008]. These latter sequences are either from Cenix bioscience and are documented in wormbase [Schindelman et al., 2011] or were designed in the framework of this project (see Table S3). JEP:vec-33 and JEP:vec-35 were designed against *ebp-1* and *ebp-1/3*, respectively, and were disclosed in [Rodriguez-Garcia et al., 2018]. JEP vectors were constructed using Gateway technology (Invitrogen). Most RNAi treatments were performed by feeding the worm with specific bacterial clones. Feeding plates were either obtained by growing a drop of bacteria mixed with IPTG at the centre of a standard NGM plate or laying bacteria on a feeding plate, i.e. an NGM plate with indicated IPTG concentration in the agar. In all cases, plates were incubated overnight [Kamath et al., 2001]. Alternatively, RNAi were obtained by injection of dsRNA in the gonads after [Timmons and Fire, 1998]. The dataset includes control embryos for the RNAi experiments, obtained by feeding with bacteria carrying the empty plasmid L4440. We did not notice any phenotype suggesting that the meiosis was impaired during these various treatments.

### 4.4 Embryos preparation and imaging

Embryos were dissected in M9 buffer and mounted on a pad (2% w/v agarose, 0.6% w/v NaCl, 4% w/v sucrose) between a slide and a coverslip. Embryos were observed at the spindle plane using a Zeiss Axio Imager upright microscope (Zeiss, Oberkochen, Germany) modified for longterm time-lapse. First, extra anti-heat and ultra-violet filters were added to the mercury lamp light path. Secondly, to decrease the bleaching and obtain optimal excitation, we used an enhanced transmission 12-nm bandpass excitation filter centred on 485 nm (AHF analysentechnik, Tübingen, Germany). We used a Plan Apochromat 100/1.45 NA (numerical aperture) oil objective. Images were acquired with an Andor iXon3 EMCCD (electron multiplying charge-coupled device) 512 *×* 512 camera (Andor, Belfast, Northern Ireland) at 33 frames per second and using Solis software. Embryos were imaged at various temperatures as reported in Suppl Table S1. To confirm the absence of phototoxicity and photodamage, we checked for normal rates of subsequent divisions [Riddle, 1997, Tinevez et al., 2012] in our imaging conditions. Images were then stored using Omero software [Li et al., 2016] and analysed from there.

### 4.5 Centrosome-tracking assay, filtering and getting spindle length

The tracking of labelled centrosomes and analysis of trajectories were performed by a custom tracking software developed using Matlab (The MathWorks) [Pecreaux et al., 2006a, Pécréaux et al., 2016]. Tracking of −20ºC methanol-fixed *γ*-tubulin labelled embryos indicated accuracy to 10 nm. Embryo orientations and centres were obtained by cross-correlation of embryo background cytoplasmic fluorescence with artificial binary images mimicking the embryo or by contour detection of the cytoplasmic membrane using background fluorescence of used dye with the help of an active contour algorithm [Pecreaux et al., 2006b]. In this work, we accurately monitor spindle elongation from about 120 seconds before anaphase onset and up to 120 seconds after, which corresponds to the most dynamic phases of spindle elongation.

To exclude the rare tracking outliers, i.e. timepoints at which the centrosome was confused with a transient bright spot briefly appearing, we compared the raw data points to median-track computed by a running median over a 3 s window. Each point at a distance larger than 3 µm was excluded. It could correspond to a displacement at 1 µm s^*−*1^ or faster, which is about one order of magnitude larger than the maximum speed observed for centrosome during elongation or when spindle breaks (targeting *cls-2* e.g.). We repeated this filtering procedure a second time, computing the median on the data points filtered in the first time. Finally, we applied quality control and ensured that no more than 1 % of the points were removed along the whole trajectory. Embryos not complying with this condition were excluded from further analysis. They corresponded to acquisition issues like poor focus. In other embryos, the remaining raw data points after filtering are used for subsequent analysis. The spindle length is computed as the Euclidean distance between the two centrosomes.

### 4.6 Quantifying KLP-7 expression level by fluorescence

The expression level of KLP-7 was assessed by quantifying fluorescence using Image J software. From the equatorial section of the zygote, at the anaphase onset, the average fluorescence intensity of the cell was measured, and the average fluorescence outside the embryo (background) was subtracted.

## 4.7 Code availability

A github repository archived on zenodo offers a Jupyter-lab Python notebook to enable the reproduction of the computation of the PCA projection and the significant figures [Le Cunff and Pecreaux, 2024]. It comes with exemplar data as comma-separated value files. To install this code, one will need an instance of conda; we used Anaconda (anaconda.org) and can install the proper environment using the provided yaml file PCA_code_published_from_history.yml.

## 5 Acknowledgements

Strains TH27, TH231, TH290, TH291, TH243 were a kind gift from Prof Anthony A. Hyman. Some strains were provided by the Caenorhabditis Genetics Center (CGC), which is funded by National Institutes of Health Office of Research Infrastructure Programs (P40 OD010440; University of Minnesota). Strain ANA019 was kindly offered by Dr Marie Delattre. Some strains were provided by NBRP, which is funded by the Japanese government. The bact-16 bacteria to perform *par-4(RNAi)* is a kind gift from Dr Anne Pacquelet. We thank Dr. Gregoire Michaux for the feeding clone library and technical support. We also thank Drs. Grégoire Michaux, Anne Pacquelet, Sébastien Huet, Marc Tramier and Olivier Dameron for discussions about the project. RRG and JP were supported by a Centre National de la Recherche Scientifique (CNRS) ATIP starting grant and La Ligue nationale contre le cancer. HB was supported by EMBO through a long-term postdoctoral fellowship (ALTF 326-2013). We also acknowledge Plan Cancer grant BIO2013-02, COST EU action BM1408 (GENiE), RTR siscom, La Ligue contre le cancer (comités d’Ille-et-Vilaine, des deux Sèvres, et du Maine-et-Loire), Rennes Métropole (AIS JP, HB and YLC), Region Bretagne (SAD AniDyn-MT grant and pRISM). Microscopy imaging was performed at the Microscopy Rennes Imaging Center, UMS 3480 CNRS/US 18 INSERM/University of Rennes 1.

## 5.1 Author contributions

Conceptualisation: YLC, JP; Experimental Data: LC, SP, XP, NS, DF, BM, RRG, ZA, AA, GDN, VC, MA, JP; Data curation: LC, SP, HB, JP; Formal analysis: YLC, JP; Funding acquisition: HB, JP, YLC; Investigation: YLC, LC, JP; Methodology: YLC, JP; Project administration: YLC, JP; Software: YLC, HB, JP; Supervision: JP; Validation: YLC, HB, CH, JP; Visualization: YLC, JP; Writing – original draft: YLC, JP; Writing – review and editing: YLC, CH, HB, JP.

