## Supplemental information for "Unveiling Inter-Embryo Variability in Spindle Length over time: towards Quantitative Phenotype Analysis"

### Supplemental information for the article "Unveiling Inter-Embryo Variability in Spindle Length over time: towards Quantitative Phenotype Analysis." by Le Cunff et al.

#### 1 Multilinear fit predicting Farhadifar features by regression on the PCA coefficients

We sought a linear relation between the manually selected features reported in [Farhadifar et al., 2015] and our features without priors. We first fitted individual embryos elongation curve using the Levenberg-Marquardt algorithm, and the equation $l(t) = l_0 + \frac{l_1}{1+\exp\left(\frac{-(t-t_0)}{\tau}\right)}$ after [Farhadifar et al., 2015]. The technical variables in this fitting-equation combine into three features: the average spindle length at early metaphase $l_0$, the final spindle length in anaphase $l_0 + l_1$, and the elongation rate $\alpha = l_1/4\tau$. We excluded the poorly fitted embryos, where the standard error over one of the four fitted variables was higher than 50% of the nominal value (16/1618 embryos) and the 14 additional embryos where fitting produced aberrant values, that was final length larger than 55 µm, the length of the embryo itself, or elongation rate larger than $1\,\mu m\,s^{-1}$.

Then, we performed a multilinear regression to estimate a linear relation between the average spindle length at early metaphase $l_0$ and coefficients 1 and 2 in PCA, namely $l_0 = a * C_1 + b * C_2 + c$, with $a$,$b$,$c$ the fitting parameters and using ordinary least squares and heteroskedastic consistent variance HC3 in estimating standard errors [MacKinnon and White, 1985] (Fig 2E) . It led to estimates $a \simeq 19.03 \pm 0.42$ (estimate ±standard error), $b \simeq -22.526 \pm 0.62$ and $c \simeq 12420 \pm 25$, while the R-squared for the model read $R^2 \simeq 0.75$.

We repeated the computation for the final spindle length in anaphase $l_0 + l_1$, and obtained estimates $a \simeq 22.04 \pm 0.65$, $b \simeq 30.71 \pm 1.06$ and $c \simeq 23040 \pm 45$, while $R^2 \simeq 0.58$ (Fig 2F).

We finally investigated the elongation rate $\alpha = l_1/4\tau$, and obtained estimates $a \simeq 0.216 \pm 0.034$, $b \simeq 1.08 \pm 0.11$ and $c \simeq 105.5 \pm 1.6$, while $R^2 \simeq 0.32$ (Fig 2G).

Overall, it suggests that our two first archetypes capture quite well the initial and, to a lower extent, final spindle length as estimated with manually-set features. Meanwhile, the elongation rate likely depends on more archetypes than the two first ones. Consistently, multilinear fits with the three coefficients modestly improve the $R^2$ for initial and final lengths while being more advantageous for the elongation rate, with values, respectively, $R_3^2 \simeq 0.87$, $R_3^2 \simeq 0.62$ and $R_3^2 \simeq 0.56$.

#### 2 Genes causing spindle shortening or limited elongation in late metaphase

We asked whether genes known to lead to a spindle shortening phenotype during late metaphase display a significantly different PCA projection from their control. We first identified *cls-2*, *tpxl-1* and *bub-1* [Ozlu et al., 2005, Lewellyn et al., 2010, Cheerambathur et al., 2013, Edwards et al., 2018, Cheeseman et al., 2005]. To extent this list to all related genes in our set, we extracted all genes which share at least 5 common phenotypes with these 3 genes. It indicated *air-2* and *klp-19*. We thus compared the PCA coefficients of individual embryos using the Mann-Whitney test (Suppl Table S6) and observed a significant difference in the third coefficient, at least in all cases.

#### 3 Supplemental methods

##### 3.1 Evaluating projection methods

In order to unveil the underlying similarities between these various dynamics in spindle elongation, we tested a range of dimension reduction methods, linear, unsupervised and extracting features (Table S4) [Jia et al., 2022]. For the sake of completeness, we also added two non-linear feature extraction algorithms, namely t-distributed stochastic neighbour embedding (t-SNE) and local linear embedding [Van der Maaten and Hinton, 2008]. For each algorithm and each corresponding projected coordinate, we computed the ratio between inner-condition variability and between-condition variability:

$$\texttt{score} = \frac{\Sigma_{k=1}^{K} n_k (\bar{x_k} - \bar{x})^2}{\Sigma_{k=1}^{K} \Sigma_{x_j \in g_k} (x_j - \bar{x_k})^2}$$

where $K$ is the number of experimental conditions in the whole dataset $g_K$, $\bar{x_K}$ the average coordinates of the group $K$, $\bar{x}$ the average over the whole dataset. We also denoted $x_j$ the coordinate of the individual embryo $j$. We obtained the global score by summing the scores obtained for each dimension. This score is analogous to the statistics used in anova analysis: the higher the score, the more separated the experimental conditions are and the more compact each group of individual experiments within the same experimental condition is in the projection space. It quantifies that experimental replicas tend to cluster together preferentially. Among linear methods that also enable us to interpret the resulting dimensions, Principal Component Analysis (PCA) performed best in clustering our dataset's various experimental conditions (Table S4). Considering PCA, and since our data include replicas of the same experiment, i.e. data points with the same label, we asked whether these are clustered by the projection methods by computing the aforementioned score upon scrambling gene labels; we repeated this assay 10000 times. It produced a Gaussian distributed histogram of mean 0.086 and standard deviation 0.01 (Fig S1D).

##### 3.2 Mapping features on the projection plan

Each experimental elongation curve is projected using PCA and is described as a point in a 2-dimensional space corresponding to its 2 first PCA coefficients. We concurrently fitted the elongation after [Farhadifar et al., 2016] and extracted the corresponding features (see main text). To detect a potential gradient of these features in the projection, we fitted a plane in the 3D space (Suppl Text § 1). Then, we extracted the corresponding 2D-gradient of such a plan and depicted it on the PCA plan as an arrow.

##### 3.3 Machine Learning

To predict the final spindle length (see section 2.2), we used a neural network consisting of seven fully connected layers of 64 neurons each, with a ReLu activation function implemented in Python with the Keras package. This network took a 25 s duration curve (spindle length) as input, i.e. about 800 data points. Alternatively, we put in the coefficients obtained by PCA projection and, in such a case, used a smaller network comprising three hidden layers of 16 neurons each. We computed the spindle length in late anaphase ($l_{LA}$), as the average of the 300 last data points spanning between 111.7 s and 120 s after anaphase onset and filtered out the embryos giving the 1% shortest and longest values. Then, we performed cross-validation and split the 1584 remaining experiments disregarding experimental conditions: 60% in the training set, 8% in the validation set and 32% in the testing set. We trained using the adam algorithm, mean squared-error loss with batch size equal to 32. We stopped after 200 epochs when using a 25 s duration spindle elongation curve as input and 150 epochs when using PCA projection of elongation over this same interval. Plotting the loss over the validation set did not suggest overfitting. We computed the predicted spindle final length and compared it to the experimental value by computing the Pearson correlation coefficient that we used as the predictive power.

##### 3.4 Bootstrapping PCA

We used a bootstrap approach to test the robustness of archetypes extracted by PCA. We set to randomly select 500 experiments among the whole dataset of 1618 embryos. It ensures sampling some but few embryos of each of the conditions (Table S1). In doing so, we measured the archetypes' robustness to changes in the dataset size and composition (Fig 4A). Because the high-frequency noise in the elongations is unimportant, we applied a running-window median filtering with size 49 points, corresponding to 1.5 s, to each track before including it in the computation. We made 500 iterations of bootstrapping.

### Supplemental figures

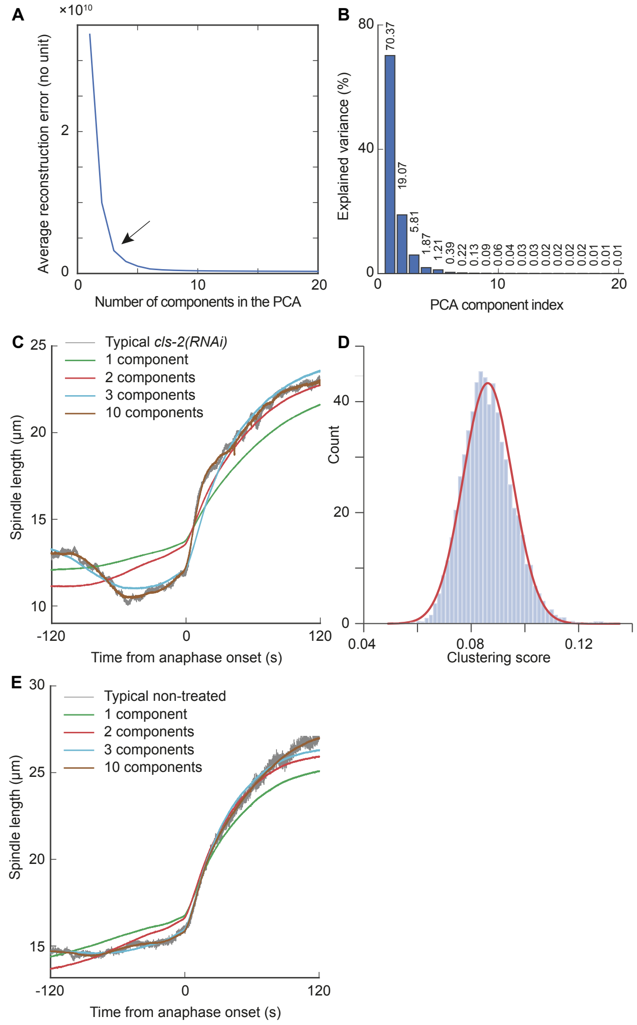


**Figure S1: Selecting the number of components for PCA projection.** (**A**) Decrease of the L2 error, i.e. the sum of the squares of the residuals, when increasing the number of dimensions in the PCA method. We suggest that 3 dimensions are optimal as it corresponds to the corner of the L-shaped curve (arrowhead). (**B**) Percentage of explained variance by each PCA component. (**C, E**) (grey) Comparing the raw spindle elongation of an exemplar single embryo labelled by γ-TUB::GFP, treated by (C) *cls-2(RNAi)* during 24 h or (E) non treated, and imaged at 18°C. (coloured curves) We reconstructed the variability around the average with 1-3 and 10 PCA components and added the average elongation of all conditions used in this paper. The third component (archetype) was essential to recapitulate the key features of the experimental trajectories, especially the transient spindle limited-elongation / shortening before anaphase onset. (**D**) Histogram of clustering scores from a PCA with scrambled labels (see suppl mat §3.1) to be compared to PCA with real labels scoring 1.86. The red line depicts the maximum likelihood Gaussian fit.

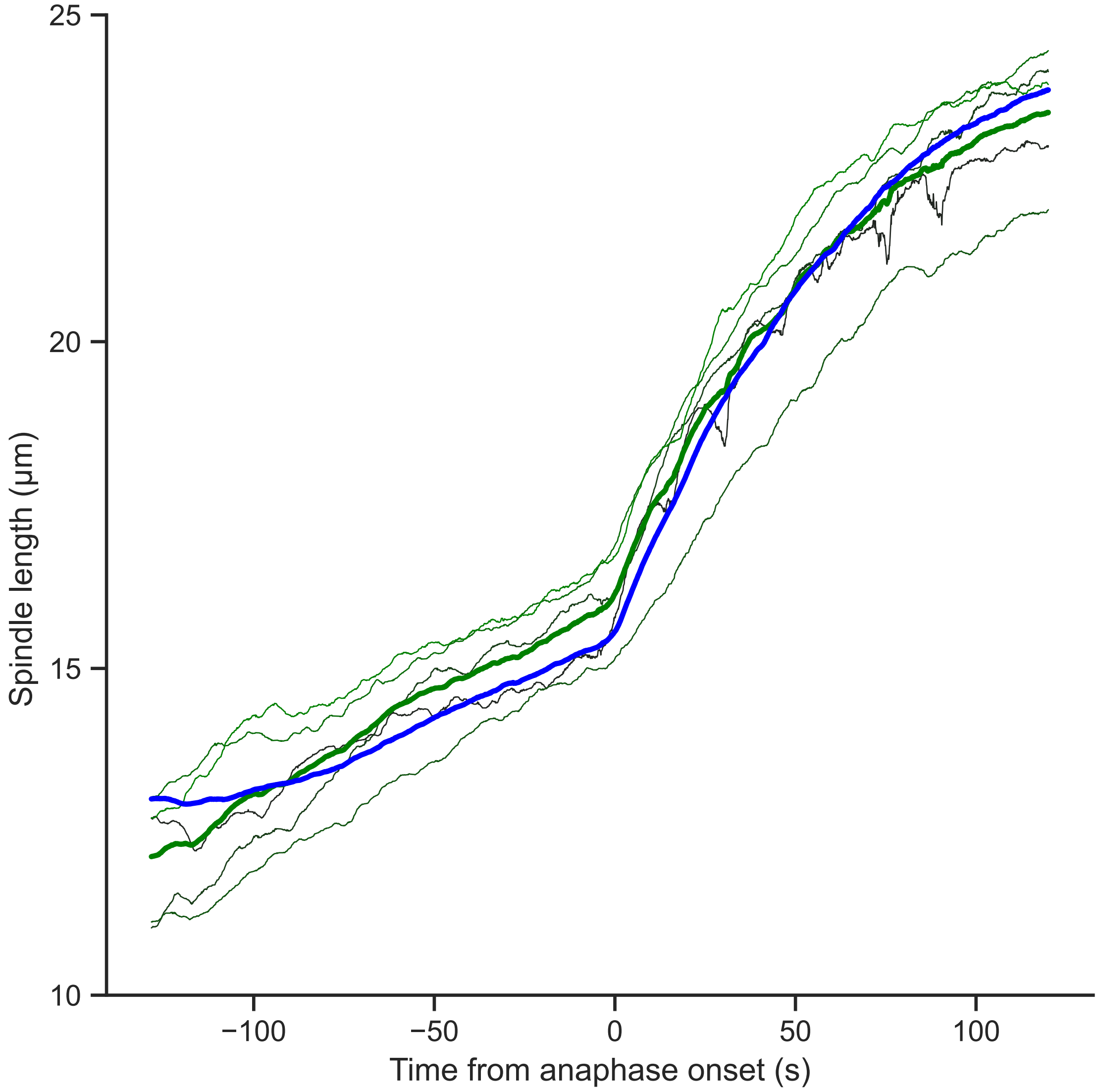


**Figure S2: Spindle elongation in non-treated embryos imaged at 18°C with low coefficient 3**. Green thin lines report individual embryo elongation curves for the $N = 5$ embryos with lowest coefficient 3, $\overline{C_3^-} \simeq -21.4 \pm 4.3$. We first selected embryos with coefficient 2 between the first and third quartiles, termed mid-coefficient 2 embryos ($N = 28$). Then among these, we took 15% of the embryos with extreme coefficient 3. Doing so, while coefficient 3 of the extreme pool is clearly different compared to the one of mid-coefficient 2 embryos, $\overline{C_3} \simeq -5.55 \pm 2.05$. In contrast, the coefficients 1 and 2 are similar in the two groups; they read $\overline{C_1^-} \simeq 53.9 \pm 16.9$ and $\overline{C_2^-} \simeq 6.8 \pm 3.7$ for the group with lowest coefficient 3 compared to $\overline{C_1} \simeq 47.2 \pm 5.02$ and $\overline{C_2} \simeq 9.73 \pm 1.23$ for mid-coefficient 2 group. The thick coloured line corresponds to the averages over these groups. The thicker blue line corresponds to the average over mid-coefficient 2 embryos. Compared to this latter average, a faster spindle elongation in late metaphase is visible in the low-coefficient-3 embryos average. All experiments were done using strain TH27, acquired at 18°C. Individual embryo and averaged tracks were smoothed using a 1.5 s-running-window median.

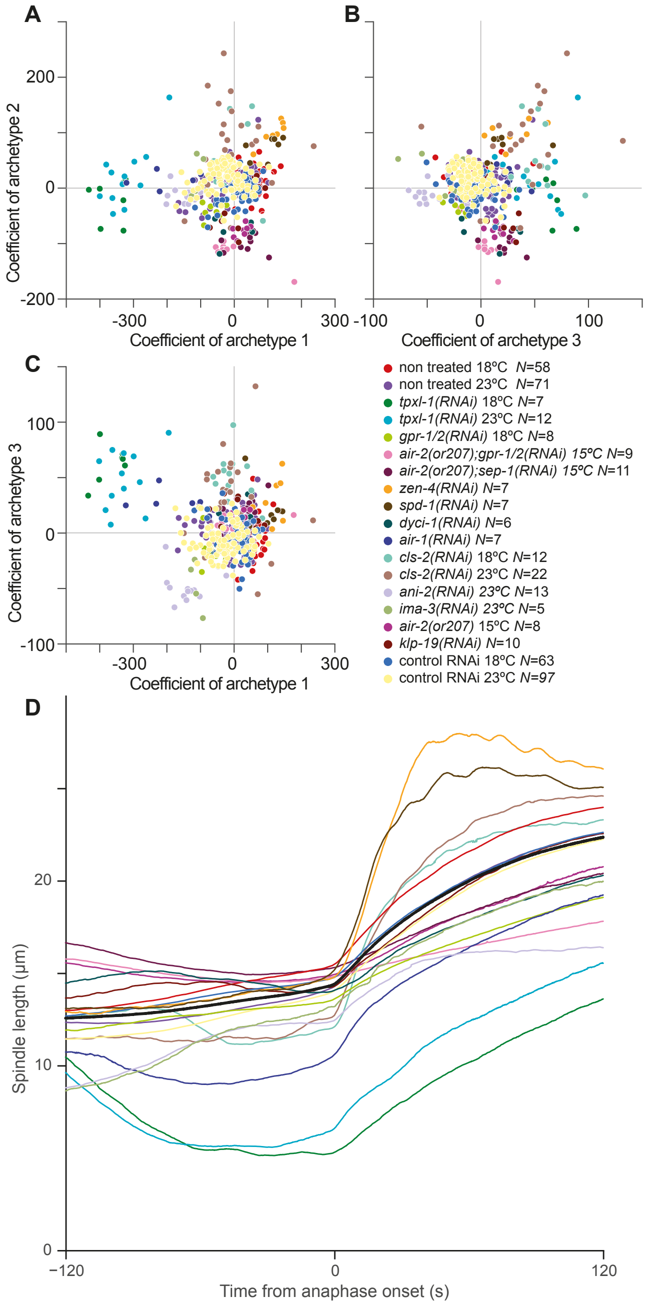


**Figure S3: Projection of the spindle elongations of individual embryos from a few conditions**, including the ones presenting extreme coefficients (Fig 2B-D) and non-treated ones. Imaging was performed at 18°C except otherwise stated. (**A**) Coefficients corresponding to the first two main archetypes (PCA components). (**B**) Similar plots for the second and third archetypes, and (**C**) for the first and third archetypes. Colours refer to genetic perturbations. Grey lines depict the 0 on each axis. All experiments were done using strain TH27 except the ones featuring *air-2(or207)*, which used JEP31. Acquisitions were performed at 18°C except otherwise stated. An interactive 3D plot is attached as Suppl File 2. (**D**) Pole-pole distance (spindle length) averaged per condition and plotted during metaphase and anaphase for the cases displayed in panels A-C. Multiple conditions treating the same gene by RNAi or mutating it are merged. Averaged tracks were smoothed using a 1.5 s-running-window median. The black thicker line corresponds to the average over the whole dataset.

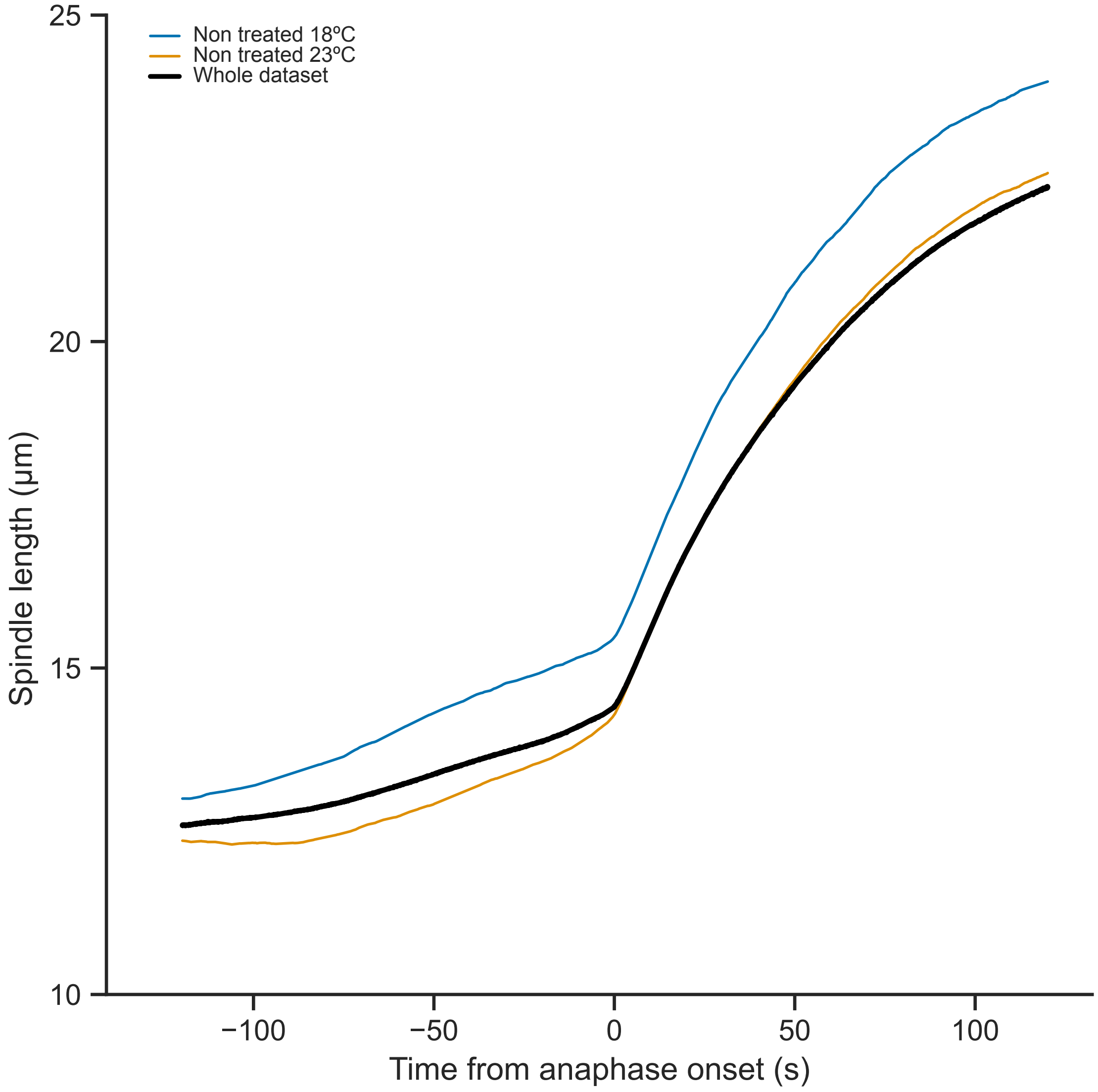


**Figure S4: Spindle elongation averaged over non-treated embryos** imaged at (blue curve) 18°C and (orange curve) 23°C. Black thicker line corresponds to the average over the whole dataset, including all conditions. All experiments were done using strain TH27. Tracks were smoothed using a 1.5 s-running-window median.

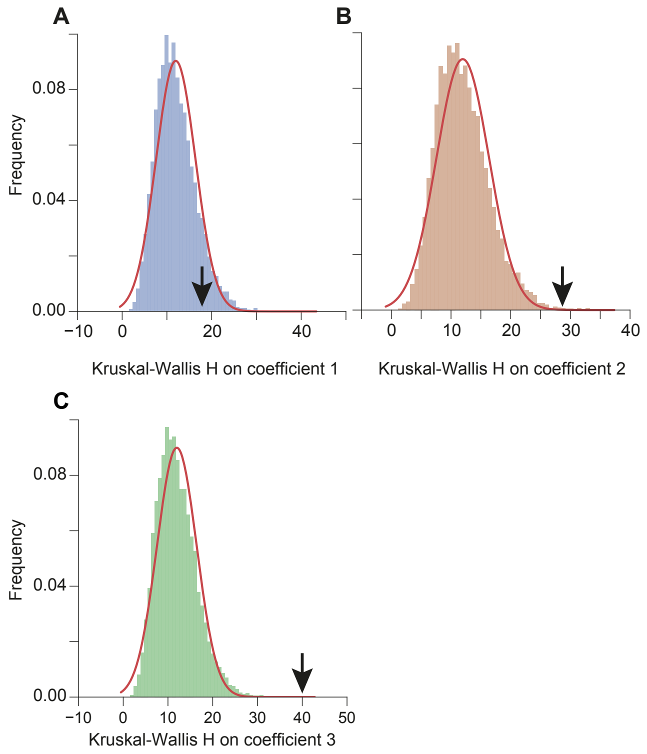


**Figure S5: Clustering of conditions per function group**. We shuffled the group labels with respect to Suppl Table S1 and computed the Kruskal-Wallis H to assess whether conditions from the same group clustered. (**A-C**) We repeated this computation 10000 times and reported the distribution of H for each coefficient. The arrows indicate the value obtained with true labels for each coefficient, $H_1 = 17.9$, $H_2 = 28.7$ and $H_3 = 39.9$. Red lines depict the maximum-likelihood Gaussian fit.

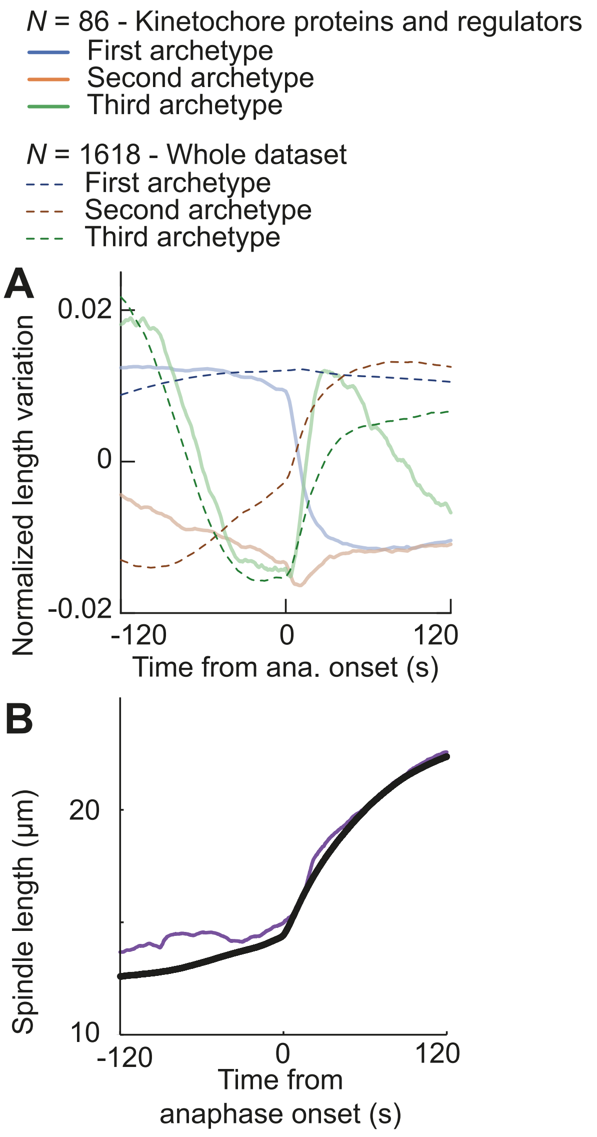


**Figure S6: Archetypes upon PCA of kinetochore functional group embryos.** (**A**) Average of the three first PCA archetypes computed considering only the embryos from conditions of the group "kinetochore proteins and regulators" (kt) ($N$=86) and compared to (dashed lines) archetypes extracted from the whole set of conditions ($N$=1618). The elongation curves were smoothed with a 1.5 s running-median filtering before computing PCA. Explained variance is reported in Table S5. (**B**) The corresponding spindle elongation was computed as the median of the average elongation curve among embryos from the same conditions. The track was smoothed using a 1.5 s-running-window median. The black thicker line corresponds to the average over the whole dataset, including all conditions.

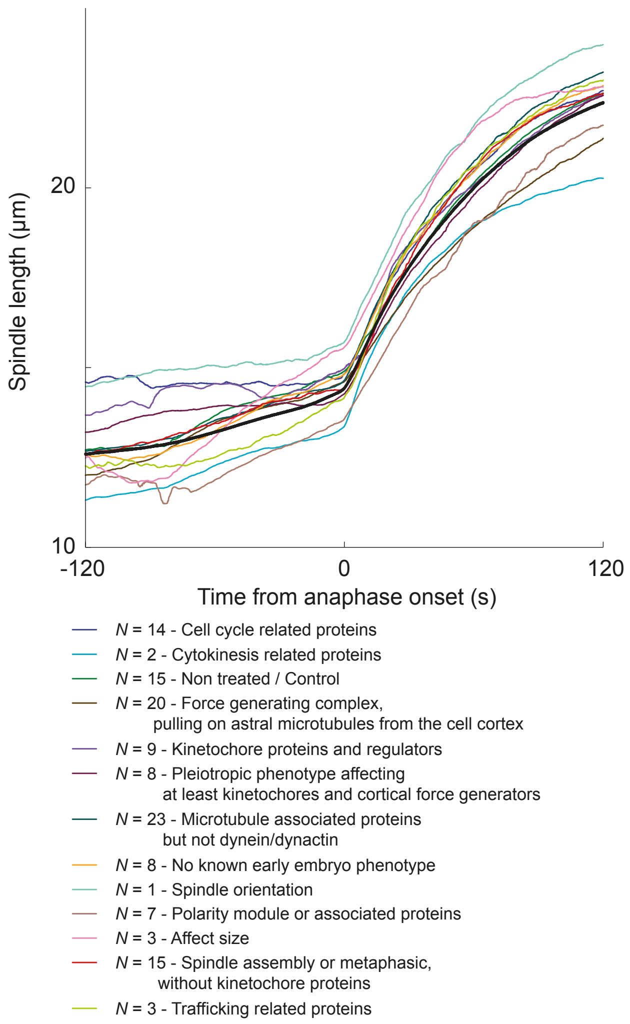


**Figure S7: Spindle elongation per group** computed as the median of the curves obtained for each condition in the group. In turn, the elongation for each condition is computed as the average of the curves for each embryo within the condition. Each group track was smoothed using a 1.5 s-running-window median. The black thicker line corresponds to the average over the whole dataset, including all conditions. The corresponding PCA values are reported at Fig 6.

#### Supplemental tables

**Table S1: Treatments used in this study.** The fluorescent strains used, carrying possibly mutated genes, are detailed in Table S2. Target identifies the condition in the figures. RNAi treatments were performed by feeding or injection as detailed in methods § 4.3. L4440 correspond to control conditions for RNAi treatments. Group corresponds to manual tags by function in the one-cell embryo based on papers and known phenotypes in wormbase [Harris et al., 2010]. Group abbreviations are detailed in Fig 6.

| Strain | Target | Treatment | Details | Duration | Temperature | Embryos Nb | Group | Reference |
|---|---|---|---|---|---|---|---|---|
| TH27 | air-1 | RNAi-V5J24 | feeding-drop | 16 | 18°C | 7 | spn | this work |
| JEP31 | air-2 cls-2 | RNAi-III4J10 mutant or207 | feeding-drop | 48 | 15°C | 8 | kt | this work |
| JEP31 | air-2 gpr-1/2 | RNAi-III5C03 mutant or207 | feeding-drop | 48 | 15°C | 9 | fgc | this work |
| JEP31 | air-2 L4440 | control RNAi mutant or207 | feeding-drop | 48 | 15°C | 6 | kt | this work |
| JEP31 | air-2 sep-1 | RNAi-I7P07 mutant or207 | feeding-1mM | 24 | 15°C | 11 | kt | this work |
| JEP31 | air-2-15C | mutant or207 | | | 15°C | 8 | kt | [Severson et al., 2000] |
| JEP31 | air-2-25C | mutant or207 | | | 25°C | 3 | kt | [Severson et al., 2000] |
| ANA019 | ANA019 none | none | | | 18°C | 7 | ctrl | this work |
| TH27 | ani-2 | RNAi II-4P19 | feeding-1mM | 48 | 23°C | 12 | ckn | [Bouvrais et al., 2018] |
| TH27 | aps-1 | RNAi V-4F01 | feeding-drop | 24 | 18°C | 12 | traf | this work |
| TH27 | aspm-1 | RNAi I-4F17 | feeding-drop | 24 | 18°C | 7 | map | this work |
| TH27 | bmk-1-18C | RNAi V-9B23 | feeding-drop | 24 | 18°C | 21 | spn | this work |
| TH27 | bmk-1-23C | RNAi V-9B23 | feeding-drop | 48 | 23°C | 9 | spn | this work |
| TH27 | C27D9.1 | RNAi II-4A13 | feeding-drop | 48 | 23°C | 6 | size | [Bouvrais et al., 2018] |
| TH27 | cdk-1 | RNAi III-5O13 | feeding-drop | 12 | 18°C | 4 | cc | this work |
| TH27 | cid-1 | RNAi III-2P5 | feeding-3mM | 48 | 18°C | 10 | size | this work |
| TH27 | clip-1-18C | RNAi JEP:vec-9 | feeding-drop | 24 | 18°C | 11 | map | this work |
| JEP32 | clip-1-23C | mutant gk470 | | | 23°C | 9 | map | this work |
| TH27 | cls-2-18C | RNAi III-4J10 | feeding-drop | 24 | 18°C | 12 | kt | this work |
| TH27 | cls-2-23C | RNAi III-4J10 | feeding-3mM | 48 | 23°C | 12 | kt | this work |
| TH27 | cls-2-23C | RNAi III-4J10 | feeding-drop | 24 | 23°C | 9 | kt | this work |
| TH27 | csnk-1 | RNAi I-5K03 | feeding-1mM | 24 | 23°C | 6 | pol | this work |
| TH27 | daf-7 | RNAi III-1C04 | feeding-drop | 24 | 18°C | 6 | nop | this work |
| TH27 | dhc-1 | RNAi I-1P04 | feeding-2mM | 20 | 18°C | 5 | kt+fgc | this work |
| TH27 | dli-1 | RNAi IV-7A05 | feeding-drop | 12 | 18°C | 8 | kt+fgc | this work |
| TH27 | dli-1 | RNAi IV-7A05 | feeding-drop | 24 | 18°C | 5 | kt+fgc | [Rodriguez-Garcia et al., 2018] |
| TH27 | dnc-1 | RNAi IV-5D24 | feeding-drop | 24 | 18°C | 16 | kt+fgc | this work |
| TH27 | dyci-1 | RNAi IV-3G8 | feeding-2mM | 16 | 18°C | 6 | kt+fgc | [Rodriguez-Garcia et al., 2018] |
| JEP27 | dyci-1 ebp-2 | RNAi IV-3G8 mutant gk756 | feeding-2mM | 16 | 18°C | 8 | kt+fgc | this work |
| JEP15 | dylt-1-18C | RNAi I-3J11 | feeding-3mM | 24 | 18°C | 11 | kt+fgc | this work |
| JEP15 | dylt-1-18C | RNAi I-3J11 | feeding-3mM | 48 | 18°C | 12 | kt+fgc | this work |

| Strain | Target | Treatment | Details | Duration | Temperature | Embryos Nb | Group | Reference |
|---|---|---|---|---|---|---|---|---|
| JEP15 | dylt-1-18C | RNAi I-3J11 | feeding-drop | 24 | 18°C | 10 | kt+fgc | this work |
| TH27 | dylt-1-20C | RNAi I-3J11 | feeding-drop | 24 | 20°C | 5 | kt+fgc | this work |
| TH27 | dyrb-1 | RNAi II-6F21 | feeding-drop | 24 | 18°C | 12 | kt+fgc | this work |
| TH27 | ebp-1 | RNAi cenix:236-e2 | feeding-2mM | 24 | 18°C | 7 | map | this work |
| TH27 | ebp-1 | RNAi cenix:236-e2 | feeding-3mM | 48 | 18°C | 11 | map | [Rodriguez-Garcia et al., 2018] |
| TH27 | ebp-1 | RNAi JEP:vec-33 | feeding-2mM | 24 | 18°C | 9 | map | this work |
| TH27 | ebp-1 | RNAi JEP:vec-33 | feeding-3mM | 48 | 18°C | 10 | map | this work |
| JEP27 | ebp-1 ebp-2 | RNAi cenix:236-e2 mutant gk756 | feeding-2mM | 24 | 18°C | 9 | map | this work |
| JEP27 | ebp-1 ebp-2 | RNAi cenix:236-e2 mutant gk756 | feeding-3mM | 48 | 18°C | 12 | map | [Rodriguez-Garcia et al., 2018] |
| JEP27 | ebp-1 ebp-2 | RNAi JEP:vec-33 mutant gk756 | feeding-2mM | 24 | 18°C | 4 | map | this work |
| JEP27 | ebp-1 ebp-2 | RNAi JEP:vec-33 mutant gk756 | feeding-3mM | 48 | 18°C | 9 | map | this work |
| JEP27 | ebp-1 ebp-2 ebp-3 | RNAi JEP:vec-35 mutant gk756 | feeding-2mM | 24 | 18°C | 9 | map | this work |
| JEP27 | ebp-1 ebp-2 ebp-3 | RNAi JEP:vec-35 mutant gk756 | feeding-3mM | 48 | 18°C | 9 | map | [Rodriguez-Garcia et al., 2018] |
| TH27 | ebp-1 ebp-3 | RNAi JEP:vec-35 | feeding-2mM | 24 | 18°C | 8 | map | this work |
| TH27 | ebp-1 ebp-3 | RNAi JEP:vec-35 | feeding-3mM | 48 | 18°C | 5 | map | [Rodriguez-Garcia et al., 2018] |
| TH27 | ebp-2-18C | RNAi II-7F10 | feeding-3mM | 48 | 18°C | 8 | map | [Rodriguez-Garcia et al., 2018] |
| TH27 | ebp-2-18C | RNAi II-7F10 | feeding-drop | 24 | 18°C | 6 | map | this work |
| TH27 | ebp-2-20C | RNAi II-7F10 | feeding-drop | 18 | 20°C | 4 | map | this work |
| JEP46 | ebp2clip1 | mutant gk756 mutant gk470 | | | 18°C | 8 | map | this work |
| TH27 | efa-6-18C | RNAi IV-6P21 | feeding-drop | 24 | 18°C | 13 | fgc | this work |
| TH27 | efa-6-23C | RNAi IV-6P21 | feeding-drop | 24 | 23°C | 4 | fgc | this work |
| TH27 | egl-5 | RNAi III-4I05 | feeding-drop | 24 | 18°C | 7 | nop | this work |
| TH27 | F21H12.2 | RNAi II-4F17 | feeding-3mM | 48 | 23°C | 7 | nop | this work |
| TH27 | goa-1 | RNAi I-3P15 | feeding-drop | 24 | 18°C | 15 | fgc | this work |
| TH65 | goa-1 gpa-16 | RNAi III-1K08 | feeding-3mM | 48 | 23°C | 9 | fgc | this work |
| TH27 | gpa-16 | RNAi II-9I-18 | feeding-drop | 24 | 18°C | 8 | fgc | this work |
| TH27 | gpb-1 | RNAi II-8A5 | feeding-1mM | 24 | 18°C | 8 | fgc | this work |
| TH27 | gpr-1 gpr-2-18C | RNAi III-4J09 RNAi III-4J09 | feeding-drop | 24 | 18°C | 8 | fgc | [Pécréaux et al., 2016] |
| TH27 | gpr-1 gpr-2-23C | RNAi III-4J09 RNAi III-4J09 | feeding-1mM | 4-6 | 23°C | 15 | fgc | [Bouvrais et al., 2018] |
| TH65 | gpr-1 gpr-2-23C | RNAi III-4J09 RNAi III-4J09 | feeding-4mM | 48 | 23°C | 7 | fgc | [Bouvrais et al., 2018] |
| TH27 | ima-3 | RNAi IV-3H04 | feeding-1mM | 34 | 23°C | 5 | size | [Bouvrais et al., 2018] |
| JEP1 | JEP1 klp-13-18C | mutant tm3737 | | | 18°C | 19 | map | this work |
| JEP13 | JEP13gpr-1 | mutant ok2126 | | | 18°C | 7 | fgc | [Pécréaux et al., 2016] |
| JEP14 | JEP14gpr-2-18C | mutant ok1179 | | | 18°C | 6 | fgc | [Pécréaux et al., 2016] |

| Strain | Target | Treatment | Details | Duration | Temperature | Embryos Nb | Group | Reference |
|---|---|---|---|---|---|---|---|---|
| JEP14 | JEP14gpr-2-23C | mutant ok1179 | | | 23°C | 6 | fgc | this work |
| JEP15 | JEP15 L4440 | control RNAi | feeding-3mM | 24 | 18°C | 11 | ctrl | this work |
| JEP15 | JEP15 L4440 | control RNAi | feeding-drop | 24 | 18°C | 7 | ctrl | this work |
| JEP15 | JEP15 none | none | | | 23°C | 6 | ctrl | [Bouvrais et al., 2018] |
| JEP27 | JEP27 none ebp-2 | none | | | 18°C | 8 | map | this work |
| JEP29 | JEP29 none 15C | none | | | 15°C | 9 | ctrl | this work |
| JEP29 | JEP29 none 25C | none | | | 25°C | 6 | ctrl | this work |
| JEP3 | JEP3 gpr-1 | mutant ok2126 | | | 18°C | 8 | fgc | this work |
| JEP4 | JEP4 gpr-2-18C | mutant tm964 | | | 18°C | 9 | fgc | this work |
| JEP4 | JEP4 gpr-2-23C | mutant tm964 | | | 23°C | 6 | fgc | this work |
| JEP5 | JEP5 mbk-2 | mutant ne992 | | | 18°C | 23 | cc | this work |
| JEP6 | JEP6 lin-5-18C | mutant ev571 | | | 18°C | 6 | fgc | this work |
| TH27 | klp-10 | RNAi IV-3N16 | feeding-drop | 24 | 18°C | 2 | map | this work |
| TH27 | klp-11 | RNAi IV-4O14 | feeding-drop | 24 | 18°C | 8 | map | this work |
| TH27 | klp-12 | RNAi IV-6G03 | feeding-drop | 24 | 18°C | 6 | map | this work |
| TH27 | klp-13-18C | RNAi X-3C14 | feeding-3mM | 24 | 18°C | 7 | map | this work |
| TH27 | klp-13-18C | RNAi X-3C14 | feeding-drop | 24 | 18°C | 6 | map | this work |
| TH27 | klp-13-23C | RNAi X-3C14 | feeding-3mM | 24 | 23°C | 11 | map | this work |
| TH27 | klp-15 | RNAi I-2F15 | feeding-drop | 24 | 18°C | 9 | spn | this work |
| TH27 | klp-16 | RNAi I-4P13 | feeding-3mM | 48 | 18°C | 9 | spn | this work |
| TH27 | klp-16 | RNAi I-4P13 | feeding-drop | 24 | 18°C | 15 | spn | this work |
| TH27 | klp-17 | RNAi II-7F24 | feeding-drop | 24 | 18°C | 3 | spn | this work |
| TH27 | klp-17 | RNAi II-7H02 | feeding-drop | 24 | 18°C | 3 | spn | this work |
| TH27 | klp-18 | RNAi IV-3O14 | feeding-drop | 24 | 18°C | 8 | spn | this work |
| TH27 | klp-19 | RNAi III-623 | feeding-drop | 24 | 18°C | 10 | kt | this work |
| TH27 | klp-20 | RNAi JEP:vec-06 | feeding-drop | 24 | 18°C | 9 | map | this work |
| TH27 | klp-3 | RNAi II-5H18 | feeding-drop | 24 | 18°C | 7 | kt | this work |
| TH27 | klp-4 | RNAi X-2K05 | feeding-drop | 24 | 18°C | 8 | map | this work |
| TH27 | klp-6 | RNAi III-2J07 | feeding-drop | 24 | 18°C | 3 | map | this work |
| TH27 | klp-7 | RNAi III-5B24 | feeding-drop | 24 | 18°C | 9 | spn | this work |
| TH27 | klp-7 | RNAi III-5B24 | feeding-drop | 36 | 18°C | 6 | spn | this work |
| TH27 | klp-7 | RNAi III-5B24 | feeding-drop | 48 | 18°C | 4 | spn | this work |
| TH27 | klp-8 | RNAi X-2I-17 | feeding-drop | 24 | 18°C | 8 | nop | this work |
| JEP27 | L4440 ebp-2 | control RNAi mutant gk756 | feeding-2mM | 24 | 18°C | 10 | map | this work |
| JEP27 | L4440 ebp-2 | control RNAi mutant gk756 | feeding-3mM | 48 | 18°C | 8 | map | this work |
| TH27 | L4440-18C | control RNAi | feeding-drop | 24 | 18°C | 13 | ctrl | this work |
| TH27 | L4440-18C | control RNAi | feeding-drop | 48 | 18°C | 6 | ctrl | [Rodriguez-Garcia et al., 2018] |
| TH27 | L4440-18C | control RNAi | feeding-1mM | 6-10 | 18°C | 10 | ctrl | this work |
| TH27 | L4440-18C | control RNAi | feeding-1mM | 6-10 | 18°C | 3 | ctrl | this work |
| TH27 | L4440-18C | control RNAi | feeding-3mM | 24 | 18°C | 5 | ctrl | this work |
| TH27 | L4440-18C | control RNAi | feeding-3mM | 48 | 18°C | 8 | ctrl | this work |
| TH27 | L4440-18C | control RNAi | feeding-drop | 24 | 18°C | 9 | ctrl | this work |
| TH27 | L4440-18C | control RNAi | feeding-drop | 24 | 18°C | 9 | ctrl | this work |
| TH27 | L4440-20C | control RNAi | feeding-drop | 24 | 20°C | 3 | ctrl | this work |

| Strain | Target | Treatment | Details | Duration | Temperature | Embryos Nb | Group | Reference |
|---|---|---|---|---|---|---|---|---|
| TH27 | L4440-23C | control RNAi | feeding-1mM | 24 | 23°C | 11 | ctrl | [Bouvrais et al., 2018] |
| TH27 | L4440-23C | control RNAi | feeding-1mM | 48 | 23°C | 4 | ctrl | this work |
| TH27 | L4440-23C | control RNAi | feeding-1mM | 6 | 23°C | 14 | ctrl | [Bouvrais et al., 2018] |
| TH27 | L4440-23C | control RNAi | feeding-3mM | 48 | 23°C | 28 | ctrl | this work |
| TH27 | L4440-23C | control RNAi | feeding-drop | 24 | 23°C | 8 | ctrl | this work |
| TH27 | L4440-23C | control RNAi | feeding-1mM | 6-10 | 23°C | 8 | ctrl | [Bouvrais et al., 2018] |
| TH27 | L4440-23C | control RNAi | feeding-4mM | 24 | 23°C | 6 | ctrl | this work |
| TH27 | L4440-23C | control RNAi | feeding-drop | 24 | 23°C | 9 | ctrl | this work |
| TH27 | L4440-23C | control RNAi | feeding-drop | 6 | 23°C | 9 | ctrl | this work |
| TH65 | L4440-23C | control RNAi | feeding-4mM | 48 | 23°C | 10 | ctrl | [Bouvrais et al., 2018] |
| TH27 | let-99 | RNAi JEP:vec-37 | feeding-3mM | 24 | 23°C | 8 | pol | [Bouvrais et al., 2018] |
| TH27 | let-99 | RNAi JEP:vec-37 | feeding-drop | 24 | 23°C | 8 | pol | this work |
| TH27 | lin-5-18C | RNAi II-5J10 | feeding-drop | 24 | 18°C | 12 | fgc | this work |
| TH27 | lin-5-18C | RNAi II-5J10 | injection | 40 | 18°C | 10 | fgc | [Pécréaux et al., 2016] |
| TH27 | lin-5-23C | RNAi II-5J10 | feeding-3mM | 16 | 23°C | 11 | fgc | this work |
| TH27 | lov-1 | RNAi II-7K23 | feeding-drop | 24 | 18°C | 8 | ctrl | this work |
| TH27 | lov-1 | RNAi II-7K23 | injection | 41 | 18°C | 10 | ctrl | [Pécréaux et al., 2016] |
| TH27 | lrg-1 | RNAi III-5G03 | feeding-drop | 24 | 18°C | 5 | nop | this work |
| TH27 | mab-5 | RNAi III-4I09 | feeding-drop | 24 | 18°C | 5 | nop | this work |
| TH27 | mbk-2 | RNAi cenix:10-e8 | feeding-drop | 24 | 18°C | 7 | cc | this work |
| TH27 | nmy-2 | RNAi I-3L24 | feeding-drop | 24 | 18°C | 7 | pol | [Pécréaux et al., 2016] |
| TH27 | none-18C | none | | | 18°C | 58 | ctrl | [Pécréaux et al., 2016] |
| TH27 | none-23C | none | | | 23°C | 71 | ctrl | [Bouvrais et al., 2018] |
| TH27 | osm-3 | RNAi III-2J07 | feeding-drop | 24 | 18°C | 6 | map | this work |
| TH65 | par-2 | RNAi III-1K08 | feeding-3mM | 48 | 23°C | 5 | pol | [Bouvrais et al., 2021] |
| TH65 | par-2 | RNAi III-1K08 | feeding-3mM | 24 | 23°C | 9 | pol | this work |
| TH27 | par-3 | RNAi III-3A01 | feeding-1mM | 24 | 23°C | 15 | pol | [Bouvrais et al., 2018] |
| TH27 | par-3 | RNAi III-3A01 | feeding-1mM | 30 | 23°C | 6 | pol | this work |
| TH27 | par-4 | RNAi bact-16 | feeding-drop | 24 | 18°C | 15 | pol | this work |
| TH27 | pkd-2 | RNAi IV-7P23 | feeding-drop | 24 | 18°C | 6 | nop | this work |
| TH27 | plk-1-18C | RNAi III-4C10 | feeding-drop | 16 | 18°C | 1 | cc | this work |
| TH27 | plk-1-18C | RNAi III-4E08 | feeding-drop | 16 | 18°C | 2 | cc | this work |
| TH27 | plk-1-23C | RNAi III-4C10 | feeding-drop | 6 | 23°C | 10 | cc | this work |
| TH27 | ptl-1 | RNAi III-1A12 | feeding-drop | 24 | 18°C | 12 | map | this work |
| TH27 | ptl-1 | RNAi III-1A24 | feeding-drop | 24 | 18°C | 13 | map | this work |
| TH27 | ptl-1 | RNAi III-1A24 | feeding-drop | 36 | 18°C | 10 | map | this work |
| TH27 | spat-1 | RNAi II-9O07 | feeding-drop | 24 | 18°C | 6 | cc | this work |
| TH27 | spd-1 | RNAi I-7D17 | feeding-drop | 24 | 18°C | 7 | spn | this work |
| TH27 | spd-2-18C | RNAi I-4O08 | feeding-1mM | 6-10 | 18°C | 19 | spn | this work |
| TH27 | spd-2-23C | RNAi I-4O08 | feeding-1mM | 6 | 23°C | 9 | spn | [Bouvrais et al., 2018] |
| TH27 | spn-4 | RNAi JEP:vec-11 | feeding-drop | 24 | 18°C | 8 | ori | this work |
| JEP16 | such-1 | mutant h1960 | | | 23°C | 11 | cc | [Bouvrais et al., 2018] |
| JEP16 | such-1 dylt-1 | mutant h1960 RNAi I-3J11 | feeding-3mM | 24 | 18°C | 9 | cc | this work |
| JEP16 | such-1 dylt-1 | mutant h1960 RNAi I-3J11 | feeding-3mM | 48 | 18°C | 10 | cc | this work |

| Strain | Target | Treatment | Details | Duration | Temperature | Embryos Nb | Group | Reference |
|---|---|---|---|---|---|---|---|---|
| JEP16 | such-1 dylt-1 | mutant h1960 RNAi I-3J11 | feeding-3mM | 96 | 18°C | 2 | cc | this work |
| JEP16 | such-1 L4440 | mutant h1960 - control RNAi | feeding-3mM | 24 | 18°C | 9 | cc | this work |
| JEP16 | such-1 L4440 | mutant h1960 - control RNAi | feeding-drop | 24 | 18°C | 8 | cc | this work |
| JEP10 | such-1 mdf-1 | mutants h1960 gk2 | | | 18°C | 22 | cc | this work |
| JEP17 | such-1 mdf-1 dylt-1 | mutants h1960 gk2 RNAi I-3J11 | feeding-3mM | 48 | 18°C | 7 | cc | this work |
| JEP17 | such-1 mdf-1 L4440 | mutants h1960 gk2 - control RNAi | feeding-3mM | 24 | 18°C | 8 | cc | this work |
| TH27 | tac-1 | RNAi II-9C20 | feeding-drop | 24 | 18°C | 7 | map | this work |
| TH102 | TH102 none | none | | | 18°C | 3 | ctrl | this work |
| TH231 | TH231 none | none | | | 18°C | 21 | ctrl | this work |
| TH65 | TH65 lin-5-23C | RNAi II-5J10 | feeding-3mM | 24 | 23°C | 9 | fgc | [Bouvrais et al., 2021] |
| TH65 | TH65 none | none | | | 23°C | 7 | ctrl | [Bouvrais et al., 2018] |
| TH65 | TH65 par-3 | RNAi III-3A01 | feeding-3mM | 48 | 23°C | 11 | pol | [Bouvrais et al., 2018] |
| TH27 | tpxl-1-18C | RNAi I-7J23 | feeding-drop | 24 | 18°C | 4 | spn | this work |
| TH27 | tpxl-1-18C | RNAi I-7L01 | feeding-drop | 24 | 18°C | 3 | spn | this work |
| TH27 | tpxl-1-23C | RNAi I-7L01 | feeding-drop | 24 | 23°C | 12 | spn | this work |
| TH27 | ubxn-2 | RNAi JEP:vec-7 | feeding-drop | 24 | 18°C | 3 | cc | this work |
| TH27 | unc-104 | RNAi II-5G20 | feeding-drop | 24 | 18°C | 7 | traf | this work |
| TH27 | unc-116 | RNAi III-4O16 | feeding-drop | 24 | 18°C | 5 | traf | this work |
| TH27 | unc-59 | RNAi I-6N04 | feeding-drop | 24 | 18°C | 13 | ckn | this work |
| TH27 | vab-8 | RNAi V-8O01 | feeding-drop | 24 | 18°C | 2 | nop | this work |
| TH27 | vab-8 | RNAi V-8O05 | feeding-drop | 24 | 18°C | 3 | nop | this work |
| TH27 | zen-4 | RNAi IV-3I07 | feeding-drop | 24 | 18°C | 7 | spn | this work |
| TH27 | zyg-11 | RNAi II-5N06 | feeding-drop | 24 | 18°C | 6 | cc | this work |
| TH27 | zyg-9 | RNAi II-6M11 | feeding-3mM | 4 | 18°C | 11 | spn | this work |
| TH27 | zyg-9 | RNAi II-6M11 | feeding-drop | 24 | 18°C | 3 | spn | [Pécréaux et al., 2016] |

**Table S2: Fluorescently tagged strains** used in this study and their detailed genotypes. Original strains are referenced by each of the crossed strains, whereas previously disclosed ones are referenced by the corresponding publication.

| **Strain** | **Genotype** | **Crossing** | **Origin and Reference** |
|---|---|---|---|
| ANA019 | C. *briggsae pie-1::Ce-tbg-1::GFP; Ce-sid-2* | | [Riche, 2015] |
| JEP1 | *unc-119(ed3) III; ddIs6 [Ppie-1::GFP::tbg-1; unc-119(+)] V; klp-13(tm3737) X* | TH27 x TM3737 | [Oegema et al., 2001, elegans Deletion Mutant Consortium, 2012] |
| JEP3 | *ddIs6 [Ppie-1::GFP::tbg-1; unc-119(+)] V; gpr-1(ok2126) III* | TH27 x VC1670 | [Oegema et al., 2001, Barstead et al., 2012] |
| JEP4 | *ddIs6 [Ppie-1::GFP::tbg-1; unc-119(+)] V; gpr-2(tm964) III* | TH27 x TM964 | [Oegema et al., 2001, elegans Deletion Mutant Consortium, 2012] |
| JEP5 | *unc-119(ed3) III; ddIs6 [Ppie-1::GFP::tbg-1; unc-119(+)] V; mbk-2(ne992) IV* | TH27 x WM73 | [Oegema et al., 2001, Pang et al., 2004] |
| JEP6 | *unc-119(ed3) III; ddIs6 [Ppie-1::GFP::tbg-1; unc-119(+)] V; lin-5(ev571) II* | TH27 x SV124 | [Oegema et al., 2001, Lorson et al., 2000] |
| JEP10 | *such-1(h1960) III; unc-46(e177) mdf-1(gk2) V. ddIs180[WRM062cF05 spd-2::2xTY1 GFP FRT 3xFlag;unc-119(+)]* | KR4012 x TH231 | [Tarailo et al., 2007, Decker et al., 2011] |
| JEP13 | *gpr-1(ok2126) III. unc-119(ed3) III (?); ddIs6 [Ppie-1::GFP::tbg-1; unc-119(+)] V* | TH27 x TH290 | [Pécréaux et al., 2016] |
| JEP14 | *gpr-2(ok1179) III. unc-119(ed3) III (?); ddIs6 [Ppie-1::GFP::tbg-1; unc-119(+)] V* | TH27 x TH291 | [Pécréaux et al., 2016] |
| JEP15 | *ddIs180[WRM062cF05 spd-2:: 2xTY1 GFP FRT 3xFlag;unc-119(+)] ltIs37 [pie-1p::mCherry::his-58 (pAA64) + unc-119(+)] IV* | JEP10 x OD56 | [Bouvrais et al., 2018] |
| JEP16 | *such-1(h1960) III; ddIs180[WRM062cF05 spd-2:: 2xTY1 GFP FRT3xFlag;unc-119(+)]; ltIs37 [pie-1p::mCherry::his-58 (pAA64) + unc-119(+)] IV* | JEP10 x OD56 | [Bouvrais et al., 2018] |
| JEP17 | *such-1(h1960) III; unc-46(e177) mdf-1(gk2) V. ddIs180[WRM062cF05 spd-2:: 2xTY1 GFP FRT 3xFlag;unc-119(+)] ltIs37 [pie-1p::mCherry::his-58 (pAA64)/unc-119(+)] IV* | JEP10 x OD56 | [Bouvrais et al., 2018] |
| JEP25 | *air-2(or207) unc-13(e51) I ddIs153[WRM064C_D03::unc-119-Nat([18578] knl-1::2xTY1wEGFP3xflag)]* | EU707 x TH243 | [Severson et al., 2000, Sarov et al., 2012] |
| JEP27 | *ebp-2(gk756) II. ddIs6 [Ppie-1::GFP::tbg-1; unc-119(+)] V* | TH27 x VC1614 | [Rodriguez-Garcia et al., 2018] |
| JEP29 | *unc-119(ed3)III; ddIs153[WRM064C_D03::unc-119-Nat([18578] knl-1::2xTY1wEGFP3xflag); ddIs180[WRM062cF05 spd-2:: 2xTY1 GFP FRT 3xFlag;unc-119(+)]* | TH231 x TH243 | [Decker et al., 2011, Sarov et al., 2012] |

| Strain | Genotype | Crossing | Origin and Reference |
|---|---|---|---|
| JEP31 | *air-2(or207) unc-13(e51) I; ddIs153[WRM064C_D03::unc-119-Nat([18578] knl-1::2xTY1wEGFP3xflag); ddIs180[WRM062cF05 spd-2:: 2xTY1 GFP FRT 3xFlag;unc-119(+)]* | JEP25 x JEP29 | [Severson et al., 2000] |
| JEP32 | *clip-1(gk470) III; ddIs6 [Ppie-1::GFP::tbg-1; unc-119(+)] V* | TH27 x VC1071 | [Rodriguez-Garcia et al., 2018] |
| JEP46 | *ebp-2(gk756) II; clip-1(gk470) III; ddIs6 [Ppie-1::GFP::tbg-1; unc-119(+)] V* | JEP27 x JEP32 | this work |
| TH27 | *unc-119(ed3) III; ddIs6 [Ppie-1::GFP::tbg-1; unc-119(+)] V* | | [Oegema et al., 2001] |
| TH65 | *unc-119(ed3); ddIs15 [pPIE-1::YFP::tba-2(genomic);unc-119(+)]* | | [Srayko et al., 2005] |
| TH102 | *N-YFP::spd-5* | | [Greenan et al., 2010] |
| TH231 | *unc-119(ed3)III; ddIs180[WRM062cF05 spd-2:: 2xTY1 GFP FRT 3xFlag;unc-119(+)]* | | [Decker et al., 2011] |
| LP447 | *klp-7(cp178[klp-7::mNG-C1^3xFlag]) III* | | [Heppert et al., 2018] |

**Table S3: Bacterial clones designed for this study** to silence genes by RNAi.

| Name | Target | Primers (forward / reverse) | Reference |
|---|---|---|---|
| JEP:vec-6 | *klp-20* | 5'-AGTACATTCCGGTGGAGCAC-3'<br>5'-TAGGCAATTGCTTTGAGCTG-3' | this work |
| JEP:vec-7 | *ubxn-2* | 5'-AAAGTGAACCGCCACCAC-3'<br>5'-CAACATTTCCCAAACGGACT-3' | this work |
| JEP:vec-9 | *clip-1* | 5'-TCCCGATGGTTCAATCAGTTT-3'<br>5'-GCATCCTCCCTTTCTTTTCA-3' | this work |
| JEP:vec-11 | *spn-4* | 5'-GAGCGACACCAACCCGCAGA-3'<br>5'-ATCTGGTCACGAAGATGATGTGGGA-3' | this work |
| JEP:vec-37 | *let-99* | 5'-CCACCAAAGGCAAG-3'<br>5'-AAGTGATCTGTTCAAAATCTTCGGA-3' | this work |

| Dimension reduction method | Score |
|---|---|
| Principal Component Analysis (PCA) | 1.86 |
| Non-metric Multidimensional scaling | 1.71 |
| Multidimensional Scaling | 1.68 |
| t-distributed stochastic neighbour embedding (t-SNE, non-linear) | 1.36 |
| Local Linear Embedding (non-linear) | 2.26 |
| Independent Component Analysis | 1.43 |
| Factor Analysis | 1.41 |
| Truncated Singular-Value Decomposition | 1.74 |
| Principal Component Analysis with scrambled labels | 0.086 |

**Table S4: Ratio of inter-group variance over intra-group variance**. We compared various projection methods by assessing their ability to cluster replicas while separating experiments corresponding to distinct treatments, using the score described in Suppl Mat §3.1. We also included some non-linear/local methods for the sake of completeness, although they will not enable the interpretability expected in our specifications. Higher scores mean that the projection method performs better. The last row corresponded to the average upon 10000 repeats of shuffling experiment labels and computing the score.

| Dataset subset | $N$ | Comp. 1 (%) | Comp. 2 (%) | Comp. 3 (%) |
|---|---|---|---|---|
| Whole dataset | 1618 | 70.63 | 19.16 | 5.83 |
| Non treated (all temperatures and strains) | 129 | 72.90 | 14.50 | 6.39 |
| Treated (all temperatures and strains, RNAi or mutant) | 1308 | 70.69 | 19.26 | 5.81 |
| Non treated at 18°C (all strains) | 58 | 52.03 | 28.31 | 10.61 |
| Non treated at 23°C (all strains) | 71 | 71.48 | 15.31 | 6.61 |
| Depleting kinetochore-related proteins (group kt) | 86 | 57.25 | 28.67 | 5.55 |

**Table S5: Explained variance for PCA over a subset of dataset**. We performed a PCA analysis on a subset of the dataset and obtained the reported percentage of explained variances (see details in main text §2.3 and Fig 4). $N$ corresponds to the number of embryos in each set.

| Gene / Target | $p$ component 1 | $p$ component 2 | $p$ component 3 |
|---|---|---|---|
| *tpxl-1(RNAi)* 18°C | $1.7 \times 10^{-9}$ | 0.50 | $3.3 \times 10^{-9}$ |
| *cls-2(RNAi)* 18°C | 0.059 | $4.6 \times 10^{-4}$ | $9.3 \times 10^{-8}$ |
| *klp-19(RNAi)* 18°C | 0.48 | 0.20. | 0.0046 |
| *air-2(or207ts)* 15°C | 0.0039 | $3.5 \times 10^{-10}$ | $5.7 \times 10^{-5}$ |

**Table S6: Mann-Whitney test comparing coefficients of genes for which depletion was previously reported as causing spindle shortening during late metaphase.** The listed treatments performed at 18°C were achieved by RNAi on the TH27 strain and compared to the corresponding L4440 treated embryos. In contrast, *air-2* was a mutant reported as temperature sensitive, although we already observed phenotype at permissive temperature. It was compared to non-treated embryos from the TH27 strain at the closest temperature. Distributions are represented at Fig S3.

### Supplemental file

#### 1 PCA coefficients averaged per condition and interactively plotted in 3D

Each experiment is projected by PCA, and then a median is computed per condition (Suppl Table S1). The resulting scatter plot is attached as an interactive plot. Corresponding 2D projections are displayed in Figure 6.

#### 2 PCA coefficients for individual embryos interactively plotted in 3D for conditions plotted in Fig S3

Individual embryos presented in figure S3 are projected by PCA. The resulting scatter plot is attached as an interactive plot. The colours are the same as in the corresponding figure.
